# Endoplasmic Reticulum Chaperone Genes Encode Effectors of Long-Term Memory

**DOI:** 10.1101/2021.10.20.465191

**Authors:** Snehajyoti Chatterjee, Ethan Bahl, Utsav Mukherjee, Emily N. Walsh, Mahesh Shivarama Shetty, Amy L. Yan, Yann Vanrobaeys, Joseph D. Lederman, K. Peter Giese, Jacob Michaelson, Ted Abel

## Abstract

The mechanisms underlying memory loss associated with Alzheimer’s disease and related dementias (ADRD) remain unclear, and no effective treatments exist. Fundamental studies have shown that a set of transcriptional regulatory proteins of the nuclear receptor 4a (Nr4a) family serve as molecular switches for long-term memory. Here, we show that Nr4a proteins regulate the transcription of a group of genes encoding chaperones that localize to the endoplasmic reticulum (ER), which function to traffic plasticity-related proteins to the cell surface during long lasting forms of synaptic plasticity and memory. Nr4a transcription factors and ER chaperones are linked to ADRD in human samples as well as mouse models, and overexpressing Nr4a1 or the ER chaperone Hspa5 ameliorates the long-term memory deficits in a tau-based mouse model of ADRD, pointing towards novel therapeutic approaches for treating memory loss. Thus, our findings establish protein folding in the ER as a novel molecular concept underlying long-term memory, providing new insights into the mechanistic basis of cognitive deficits in dementia.

**One-Sentence Summary:** Molecular approaches establish protein folding in the endoplasmic reticulum as a novel molecular concept underlying synaptic plasticity and memory, serving as a switch to regulate protein folding and trafficking, and driving cognitive deficits in neurodegenerative disorders.

## Introduction

Impaired memory consolidation and the resulting long-term memory loss is an early symptom of Alzheimer’s disease and related dementias (ADRD)(*1–3*). Memory consolidation requires the transcription of new genes, in sophisticated spatial and temporal patterns, under the control of specific families of transcription factors (*4–7*). The largest class of transcription regulators in metazoans is composed of the nuclear receptor superfamily (*8*), which regulates diverse biological processes ranging from metabolism and reproduction to development and neuronal function. Among the several subclasses of nuclear receptors, the Nr4a subfamily (Nr4a1 (NUR77, NGF-IB), Nr4a2 (NURR1/HZF-3/NOT/RNR1), and Nr4a3 (NOR1)) has emerged as a critical mediator of long-term memory (*4, 5*). Notably, these ligand-independent “orphan” receptors are robustly upregulated in the hippocampus within minutes after learning (*3, 4*). The learning-dependent expression of the Nr4a genes is regulated by histone acetylation, which is driven by recruitment of cAMP-response element binding (CREB) binding protein (CBP) (*9*) to CREB response elements in the promoters of these genes (*10, 11*). Blocking the expression or inactivating the transactivation function of Nr4a factors is sufficient to impair long-term memory (*4, 5*) and synaptic plasticity (*12*), whereas transgenic or pharmacological activation enhances long-term memory (*13, 14*). Moreover, Nr4a function is impaired in brain disorders characterized by debilitating cognitive impairment, ranging from schizophrenia, Parkinson’s disease to ADRD (*3, 4, 15, 16*). However, despite the critical importance of the Nr4a subfamily, the effector genes that these transcription factors regulate in the hippocampus during memory consolidation have remained elusive. Here, we identify ER chaperone genes as downstream effector genes regulated by these transcription factors, and we establish a role for chaperone function in long-term memory and synaptic plasticity. We further demonstrate that Nr4a transcription factors and the downstream ER chaperones that they regulate are key mediators of the long-term memory loss associated with ADRD, providing new candidate targets for the development of novel therapeutic interventions.

## Results

### Nr4a transcription factors regulate expression of genes encoding ER chaperones during memory consolidation

To identify effector genes regulated by the Nr4a subfamily during memory consolidation, we used transgenic mice that express a dominant-negative form of Nr4a1 (CaMKIIα-tTA TetO-Nr4ADN) in excitatory neurons, such that the transcriptional activity of all three Nr4a family members is blocked in these cells (*4*). To assess hippocampus-dependent memory, we examined the performance of Nr4ADN mice in spatial object recognition (SOR), a task that depends on the preference of mice to explore a spatially displaced object (*17*). In a 24 hr test of long-term memory, the control mice (CaMKIIα-tTA), but not Nr4ADN mice (double transgenic: CaMKIIα-tTA, TetO-Nr4ADN), exhibited a significant preference for the displaced object (**Fig. 1A-B**). In contrast, in a 1 hr test of short-term memory, both the Nr4ADN and control mice showed a preference for the displaced object (**fig. S1**). Thus, Nr4ADN mice exhibit selective deficits in long-term spatial memory. To identify genes regulated by Nr4a transcription factors during memory consolidation, we trained Nr4ADN and control littermates in the SOR task and extracted total RNA from the dorsal hippocampus 2 hr after training (**Fig. 1C**). We chose this time point to identify effector genes targeted by the Nr4a subfamily of transcription factors, which are immediate early genes induced within minutes after training. This analysis revealed 54 differentially expressed genes (DEGs) (**Fig. 1D**, **table S1**) in Nr4ADN versus control mice, with 40 downregulated and 14 upregulated genes in Nr4ADN mice after learning (**Fig. 1D**).

**Figure 1.**
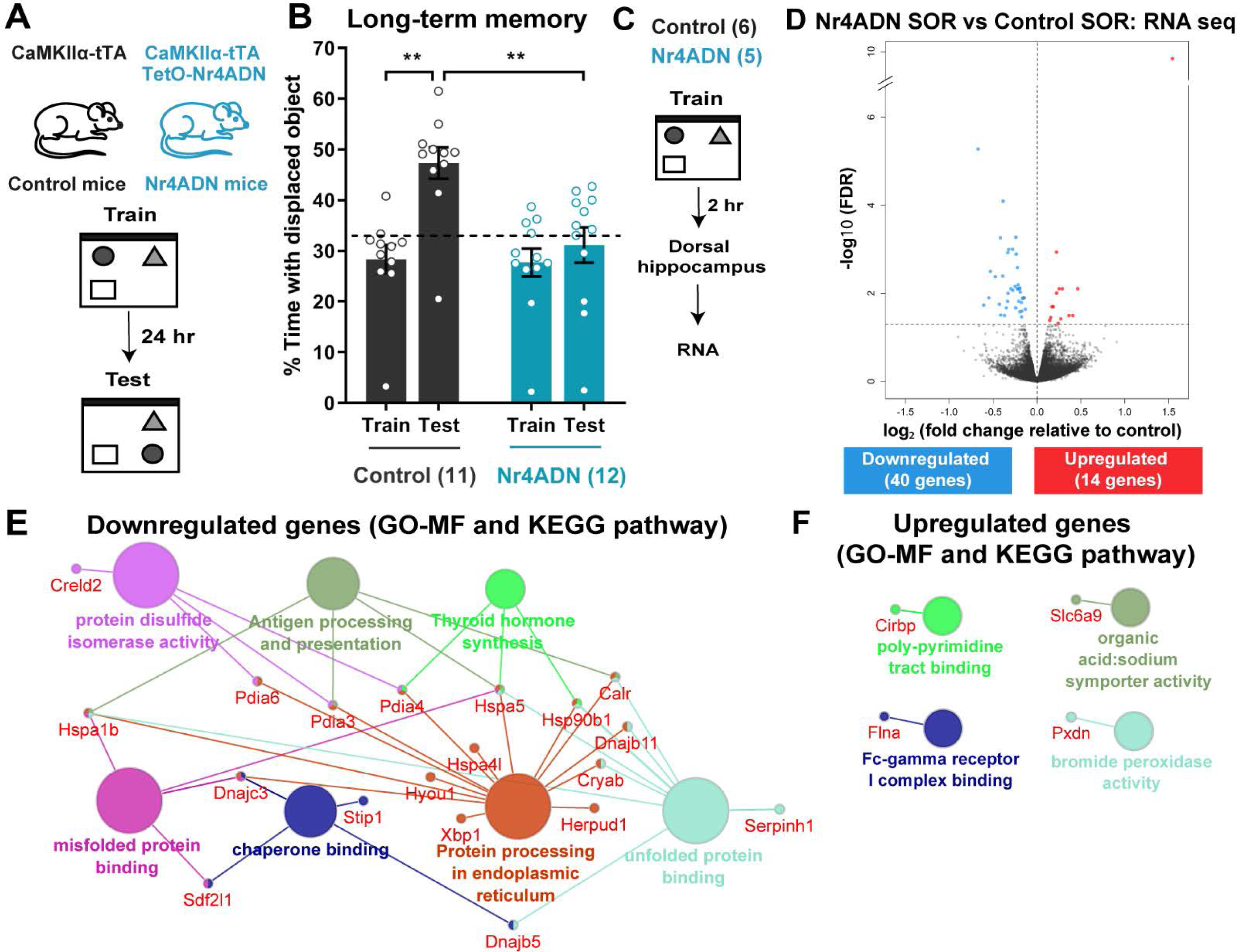
A multiprotein ER chaperone complex is downstream of Nr4a transcription factors during memory consolidation. **(A)** Schematic depicting spatial object recognition (SOR) procedure. Mice expressing the tetracycline transactivator (tTA) protein under the CaMKIIα promoter (CaMKIIα-tTA: control mice) and counterparts additionally expressing both CaMKIIα-tTA and the dominant-negative mutant Nr4A1 under the control of TetO promoter (CaMKIIα-tTA, TetO-Nr4A dominant negative: Nr4ADN mice) were trained in SOR and then tested after 24 hr. **(B)** Preference for the displaced object (DO, dotted line marks 33% chance) during the 24-hr test session relative to training. Two-way ANOVA: significant main effect of genotype (F_(1, 21)_ = 18.42, p=0.0003) and significant main effect of sessions (F_(1, 21)_ = 8.417, p=0.0085). Sidak’s multiple comparisons test: **p=0.0054 (control mice, Train versus 24-hr Test), **p= 0.0011 (control 24-hr Test versus Nr4ADN 24-hr Test). Control: n=11 (3F) Nr4ADN: 12 (4F). **(C)** Schematic depiction of RNA-seq experiment. Nr4ADN and control male mice were trained in SOR and euthanized 2 hr after the final training. mRNA was harvested from the dorsal hippocampus and processed for the preparation of an RNA-seq library. **(D),** Volcano plot illustrating significance (y-axis) and magnitude (x-axis) of the downregulation (blue) and upregulation (red) of genes in Nr4ADN mice. **(E-F),** Functional groupings of network of enriched categories for genes whose differential expression (**E**, downregulation; **F**, upregulation) was significant, using the ClueGO and CluePedia plugins of the Cytoscape software. Gene Ontology terms include Molecular Functions (MF) and Kyoto Encyclopedia of Genes and Genomes (KEGG) and are represented as nodes (κ score level ≥0.4), with node size representing the significance of the term enrichment. Only the most significant term in each group is presented in bold.

Enrichment network analysis was used to identify the pathways most represented among the down- and upregulated genes. The downregulated pathways included protein processing in the ER, chaperone binding, protein disulfide isomerase activity and several other pathways related to protein folding in the ER (**Fig. 1E**). The upregulated pathways included the poly-pyramidine tract binding pathway linked to *Cirbp* expression, a RNA binding protein associated with translational control (**Fig. 1F**). Protein-protein interaction analysis of the downregulated genes in Nr4ADN mice identified a significant cluster composed of nine chaperone proteins (Hspa5, Hsp90b1, Pdia3, Pdia4, Pdia6, Sdf2l1, Dnajb11, Hyou1, and Calr, **fig. S2**). All of these are components of a large ER multiprotein chaperone complex known to bind nascent proteins (*18*).

Next, we performed RNA-seq using the dorsal hippocampus from control mice trained in SOR (SOR training + 2 hr) or untrained mice (homecage, HC). RNA-seq analysis revealed that learning increased the expression of 42 genes and reduced expression of 9 genes (**fig. S3, table S2)**. Comparison of this gene expression data with the data from control and Nr4ADN mice after learning (**Fig. 1D**) identified 15 genes induced in control mice after learning that were downregulated in Nr4ADN mice. These genes included ER chaperone genes *Hspa5, Pdia4, Pdia6, Sdf2l1,* and *Dnajb11* (**Fig. 2A**). We next analyzed Nr4a1 occupancy on two of these candidate genes that are critical for protein folding (*Hspa5* and *Pdia6*)(*18*) using data from a previously published Nr4a1 chromatin immunoprecipitation followed by sequencing (ChIP-seq) study (*19*). The promoters of the *Hspa5* and *Pdia6* genes were found to be enriched for Nr4a1 binding motifs (**fig. S4**), suggesting that this transcription factor directly regulates the expression of these genes. We further examined the expression profiles of these two candidate genes during the first 4 hr after learning in wild-type (C57BL/6J) mice revealing that spatial learning induced the expression of both *Hspa5* and *Pdia6* genes (**Fig. 2B-C**). Subsequent investigation of these two candidate genes in Nr4ADN mice confirmed that their regulation by the Nr4a proteins occurs only after learning (**Fig. 2D-E**, **fig. S5**). Such regulation was not observed when expression of the Nr4ADN transgene was suppressed by treatment of the mice with doxycycline (Dox; **fig. S6**). These convergent data demonstrate that Nr4a transcription factors regulate the expression of a discrete set of ER chaperone genes during memory consolidation. Even though these ER chaperones are known regulators of ER stress, Nr4ADN mice do not exhibit elevated levels of key ER stress markers (p-IRE1, ATF4 and ATF6) following learning **(fig. S7**), and we see that only a subset of genes linked to the unfolded protein response is regulated by Nr4a factors following learning.

**Figure 2.**
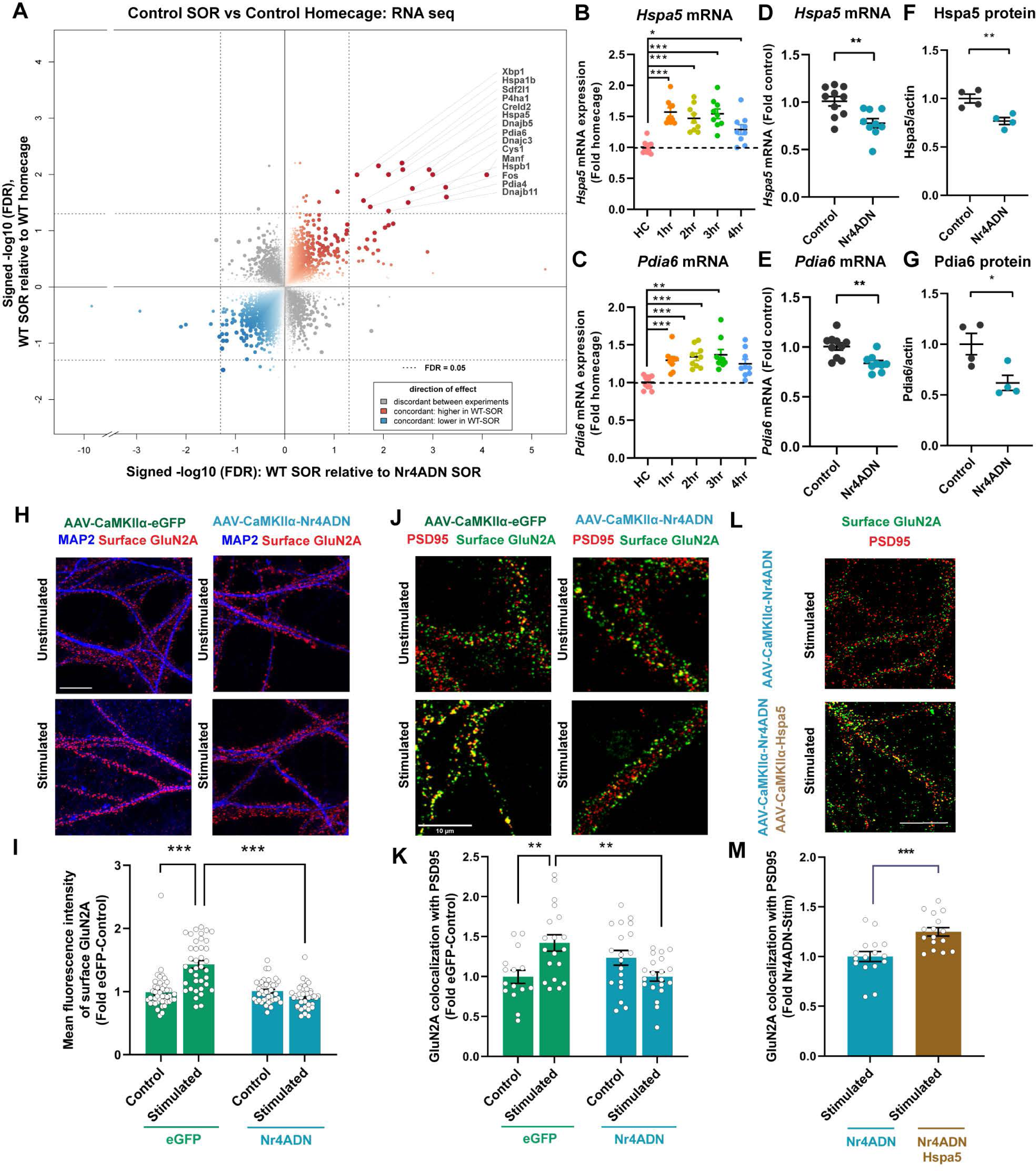
A subset of the genes downstream of Nr4a are induced by learning. **(A)** Quadrant plot based on total RNA seq data. Genes induced by learning were identified based on comparison of genes regulated in dorsal hippocampus of control male mice 2 hr after SOR (tTA^+^ Nr4ADN^-^ n=2, tTA^-^ Nr4ADN^-^ n=2) and homecage control mice (tTA^+^ Nr4ADN^-^ n=2, tTA^-^ Nr4ADN^-^ n=2). Quadrant plot comparing genes regulated by learning in control mice experiment to genes regulated by Nr4ADN after learning. Induction of genes normally upregulated by SOR is downregulated in Nr4ADN mice (labeled points). Size, opacity, and color intensity of each point reflect the minimum FDR value for a gene between each experiment. **(B-C)** Expression of the (**B**) *Hspa5* and (**C**) *Pdia6* mRNAs in C57BL/6J male mice trained in SOR and euthanized at the indicated times after final training trial (1 hr: n=9; 2 hr: n=9; 3 hr: n=9; and 4 hr: n=9), expressed as fold difference from that in mice handled only in the homecage (baseline controls, n=10). One-way ANOVA: *Hspa5*: F_(4, 41)_ =13.00, p<0.0001, Sidak’s multiple comparisons tests: ***p<0.0001 (HC versus 1 hr), ***p<0.0001 (HC versus 2 hr), ***p<0.0001 (HC versus 3 hr), *p=0.0158 (HC versus 4 hr), *Pdia6*: F_(4, 41)_ =8.442, p<0.0001, Sidak’s multiple comparisons tests: ***p=0.0008 (HC versus 1 hr), ***p=0.0001 (HC versus 2 hr), ***p<0.0001 (HC versus 3 hr), **p=0.0054 (HC versus 4 hr). **(D-E)**, Downregulation of gene expression at 2 hr after SOR training, in male Nr4ADN (n=9) and control (n=10) mice, as validated by qPCR. Unpaired t-test: t_(17)_ =3.305, **p=0.0042 (*Hspa5*), t_(17)_ =3.630, **p=0.0021 (*Pdia6*). **(F-G),** Quantification of Western blot of lysates of synaptosomes isolated from the dorsal hippocampus of male Nr4ADN (n=4) and control mice 2 hr after SOR training (n=4). Unpaired t-test: t_(6)_ =4.011, **p=0.0070 (Hspa5), t_(6)_ =2.982, *p= 0.0246 (Pdia6). **(H-K),** Cultured neurons were transduced with eGFP or Nr4ADN on DIV 16 or 17 and stimulated with KCl before live staining. **(H)** Co-staining for dendrites (MAP2 antibody) and surface GluN2A by immunofluorescence (IF) in transduced cells following KCl stimulation. Scale bar: 10 µm. **(I)** Quantification of surface staining for GluN2A in **H**, as mean fluorescence intensity. Mixed effect analysis: significant AAV construct x treatment interaction F_(1, 157)_ =38.25, p<0.0001. Sidak’s multiple comparisons test: ***p<0.0001 (control eGFP versus stimulated eGFP), ***p<0.0001 (stimulated eGFP versus stimulated Nr4ADN). **(J)** Co-staining for surface GluN2A and PSD95, by IF in transduced cells following KCl stimulation. Scale bar: 10 µm. **(K)** Quantification of surface GluN2A and PSD95 co-localization. Mixed effect analysis: significant AAV construct x treatment interaction: F_(1, 69)_ =14.57, p=0.0003. Sidak’s multiple comparison tests: **p=0.0026 (control eGFP versus stimulated eGFP), **p=0.0012 (stimulated eGFP versus stimulated Nr4ADN). **(L)** Cultured neurons were transduced with Nr4ADN or Nr4ADN+Hspa5 on DIV 16 or 17 and stimulated with KCl before live staining. Co-staining for surface GluN2A and PSD95, by IF in transduced cells following KCl stimulation. Scale bar: 10 µm. **(M)** Quantification of surface GluN2A and PSD95 co-localization. Unpaired t-test: t_(30)_ =3.727, ***p=0.0008.

Chaperones, such as Hspa5 and Pdia6, are found in synaptosomes, where they facilitate the folding and assembly of nascent polypeptides, as well as the trafficking of proteins to the neuronal surface (*20, 21*). The synaptic abundance of Hspa5 and Pdia6 was significantly lower after SOR training in Nr4ADN mice compared to controls (**Fig. 2F-G, fig. S8**). Previous studies have demonstrated that Hspa5 plays a critical role in regulating the postsynaptic membrane delivery of the N-methyl-D-aspartate (NMDA) receptor subunit GluN2A in response to neuronal stimulation (*20*). Therefore, we next investigated whether Nr4a factors might regulate the surface expression of GluN2A. We expressed Nr4ADN (or eGFP as a control) in primary hippocampal neurons using a viral-based approach and assessed the distribution of GluN2A receptors following KCl-mediated neuronal depolarization. We found significant increases in surface levels of GluN2A in eGFP-transduced cells after neuronal depolarization (**Fig. 2H-I**), whereas Nr4ADN-transduced cells failed to exhibit activity-induced trafficking of GluN2A (**Fig. 2H-I**). We also found that Nr4ADN-expressing neurons show a significant decrease in activity-induced post-synaptic surface localization of GluN2A, as evidenced by the reduced co-localization of GluN2A with the post-synaptic density protein PSD95 (**Fig. 2J-K**). Next, we investigated whether rescue of Hspa5 expression would be sufficient to increase post-synaptic localization of GluN2A in Nr4ADN-expressing neurons. Overexpression of Hspa5 increased PSD95 co-localization with GluN2A in Nr4ADN-expressing neurons (**Fig. 2L-M**). Our findings demonstrate that the regulation of chaperone protein gene expression, such as *Hspa5*, by Nr4a transcription factors is critical for the folding and synaptic trafficking of receptor proteins that are key to synaptic plasticity.

### Restoring protein chaperone function prevents long-term memory and synaptic plasticity deficits in Nr4ADN mice

Chaperones fold nascent proteins into functional three-dimensional conformations (*22*), and their upregulation after learning (**Fig. 2A-C**) is essential for the activity-dependent processing and trafficking of key synaptic proteins (**Fig. 2H-M**). Given the roles of chaperones in protein folding, we next performed experiments to determine whether the deficits in memory and synaptic plasticity observed in Nr4ADN mice are related to impairment of this process. First, we examined the effect of phenylbutyrate (PBA), a hydrophobic chemical chaperone, in Nr4ADN mice. PBA interacts with the exposed hydrophobic regions of nascent proteins to facilitate folding (*23*), partially reverses the mis-localization of proteins (*24*), and facilitates delivery of proteins that are critical for neuronal plasticity to the cell surface (*25–27*). Importantly, PBA has shown promise in rescuing cognitive impairment in several mouse models of neurodegenerative diseases (*28–31*), and these neuroprotective effects have been attributed to its chaperone activity (*28, 32*). Notably, systemic delivery of a single dose of PBA to Nr4ADN mice immediately following SOR training rescued long-term memory deficits (**Fig. 3A**); the same treatment of control mice did not augment long-term memory (**Fig. 3B**). Because PBA functions as both a molecular chaperone and an inhibitor of histone deacetylases (HDACs), we confirmed that this rescue was not due to changes in expression of Hspa5 and Pdia6 in Nr4ADN mice (**fig. S9**). Additionally, sodium butyrate (NaBu), an HDAC inhibitor that has no molecular chaperone activity (*28*), failed to rescue memory in Nr4ADN mice (**Fig. 3A**). This finding is consistent with our previous observation that broad HDAC inhibition is not sufficient to reverse the memory deficits in Nr4ADN mice (*4*).

**Figure 3.**
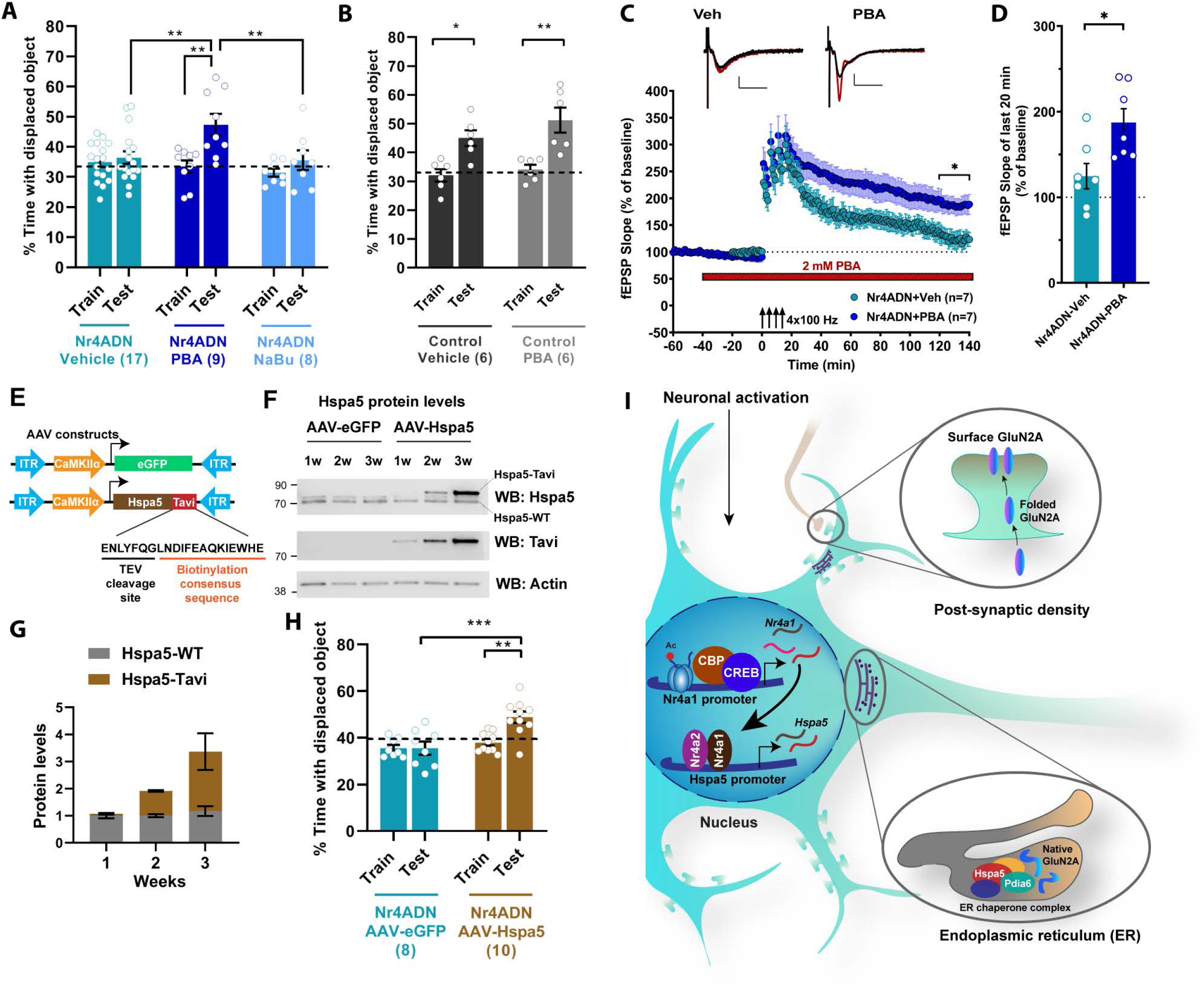
The Nr4a proteins contribute to memory through downstream chaperone proteins. **(A)** Nr4ADN mice were *i.p.* injected with phenylbutyrate (PBA, 200mg/kg, n=9), sodium butyrate (NaBu, 200mg/kg, n=8 (2F)) or vehicle (n=17 (2F)) immediately after SOR training and tested for long-term memory 24 hr later. Two-way ANOVA: significant treatment x sessions interaction (F_(2, 31)_ =4.207, p=0.0242), Sidak’s multiple comparisons tests: **p=0.0011 (Nr4ADN mice-PBA, train versus 24-hr test), **p=0.0036 (Nr4ADN-PBA 24 hr test versus Nr4ADN-Vehicle 24 hr test) and **p=0.0084 (Nr4ADN-PBA 24 hr test versus Nr4ADN-NaBu 24 hr test). **(B)** Male control mice were *i.p.* injected with PBA, (200mg/kg, n=6) or vehicle (n=6) immediately after completion of SOR training and tested for long-term memory 24 hr later. Two-way ANOVA: significant main effect of sessions F_(1, 10)_ =33.46, p=0.0002, Sidak’s multiple comparisons tests: *p=0.0110 (control mice-Vehicle, Train versus Test), **p=0.0018 (control mice-PBA, train versus test). **(C-D),** Effects of PBA on persistence of LTP, as demonstrated by representative fEPSP slope over final 20 min of recordings. Expression of Nr4ADN attenuates persistence of LTP in hippocampal slices (Nr4ADN-veh), while bath-treatment with 2 mM PBA rescues these LTP deficits (Nr4ADN-PBA) (Two-way repeated measures ANOVA, effect of PBA treatment F_(1, 12)_ = 8.125, p = 0.0146). The mean fEPSP slope over the last 20 min of the recordings was enhanced in PBA-treated slices compared to vehicle-treated slices (PBA-treated: 187.3 ± 16.2%, n = 7 slices, 4 mice; vehicle-treated: 124.6 ± 14.9%, n = 7 slices, 5 mice; Unpaired t-test, \**p* = 0.0146). Treatment with 2 mM PBA had no significant effect on the baseline responses (Pre-drug baseline, 20 min: 100.1 ± 0.13%; post-drug pre-induction baseline, 20 min: 90.63 ± 6.6%, Paired t-test, p =0.2094). The representative fEPSP traces shown are sampled at baseline (black) and at the end of the recording (red). Scale bar 2 mV, 10 ms. Error bars indicate SEM. **(E)** Schematic of viral constructs used to drive expression of Hspa5 following infusion into dorsal hippocampus of male C57BL/6J mice. AAV_9_-CaMKIIα-eGFP served as vector control and AAV_9_-CaMKIIα-Hspa5-Tavi was used to drive expression of Hspa5 in excitatory neurons. The Tavi-tag can be identified by an antibody against a consensus biotinylation sequence and has a TEV sequence that can be used to cleave it from Hspa5. **(F)** Western blot of synaptosomes, showing mild Hspa5-Tavi expression within one week of infusion, and expression approximately equal to that of endogenous Hspa5 within 2 weeks of infusion. **(G)** Quantitation of data in **F**. **(H)** Long-term memory (24 hr) assessment of Nr4ADN mice infused with AAV-eGFP or AAV-Hspa5-Tavi into dorsal hippocampus. Two-way ANOVA: significant AAV type x session interaction: F_(1, 16)_ = 6.985, p=0.0177, Sidak’s multiple comparisons tests: **p= 0.0022 (AAV-Hspa5, Train versus Test), ***p= 0.0002 (AAV-Hspa5, 24h-test versus AAV-eGFP, 24-hr test) while eGFP infused Nr4ADN mice showed no preference towards the DO. AAV-Hspa5: n=10 (4F) and AAV-eGFP: n=8 (3F). **(I)** Schematic illustration of model wherein learning-induced expression of Nr4a1 drives Hspa5 expression in nucleus to initiate protein folding in the ER that enables the expression of functional proteins at the synapse surface.

We previously showed that Nr4ADN mice exhibit deficits in a form of persistent, protein synthesis-dependent long-term potentiation (LTP) induced by repeated spaced high-frequency stimulation of the hippocampal CA1-Schaffer collateral synapses (*12*). Given that PBA treatment reversed long-term memory deficits in Nr4ADN mice, we examined its effects on this long-lasting form of LTP. Following 20 minutes of stable baseline field-excitatory postsynaptic potentials (EPSPs) recordings, hippocampal slices from Nr4ADN mice were treated by bath application of 2 mM PBA (dissolved in the artificial cerebrospinal fluid, aCSF). After 40 min of PBA treatment, long-lasting LTP was induced using a spaced 4-train stimulation protocol (four 100 Hz, 1-sec trains separated by 5 min). Potentiation in slices from Nr4ADN mice decayed quickly, thus showing deficits in the persistence of LTP, as reported in our previous study (*12*). Treatment with PBA rescued the deficits in long-lasting LTP in Nr4ADN slices leading to persistently enhanced potentiation compared to the vehicle group (**Fig. 3C-D**). At the concentration used, PBA did not have significant effects on the pre-induction baseline (**Fig. 3C**). These findings demonstrate that PBA treatment reverses the deficits in long-lasting synaptic plasticity and memory in Nr4ADN mice by promoting the folding of newly synthesized proteins.

To define the specific role of the molecular chaperone Hspa5 in the memory deficits observed in Nr4ADN mice, we reinstated Hspa5 expression selectively in hippocampal excitatory neurons using a viral approach and performed behavioral studies two weeks after viral infusion (**Fig. 3E-G**). The level of overexpression achieved was sufficient to reverse the long-term spatial memory deficits observed in Nr4ADN mice (**Fig. 3H**), supporting the idea that Hspa5 is downstream of Nr4a transcription factors (**Fig. 3I**). These findings suggest that the deficits in synaptic plasticity and long-term memory in Nr4ADN mice are due to disruption of a chaperone activity required for the folding of newly synthesized proteins. Overall, we conclude that the activity induced regulation of ER chaperone Hspa5 by Nr4a transcription factors is essential for the native protein folding required for consolidation of long-term memory.

### Activation of Nr4a1 or ER chaperone function ameliorates memory impairments in an ADRD mouse model

Nr4a transcription factors were previously implicated in Aβ aggregation and memory deficits (*33*). Therefore, we investigated the expression of Nr4a transcripts in the hippocampus of ADRD patients with increasing grades of pathology. Using a database of RNA sequencing and pathological findings from post-mortem brain tissue from patients and healthy controls in The Allen Brain Institute study of Aging, Dementia, and Traumatic Brain Injury (*34*), we examined the relationship between hippocampal expression of *NR4A* subfamily members across increasing Cerad (Consortium to Establish a Registry for Alzheimer’s Disease) score, a neuropsychological assessment of the progression of AD, and Braak stage, a measure of the distribution and pathological burden of neurofibrillary tangles (NFTs). We found that levels of expression of *NR4A1* and *NR4A2* were negatively correlated with both the Cerad scores (**Fig. 4A**) and Braak stages (**Fig. 4B**), both measures of the severity of ADRD pathology, whereas NR4A3 was not significantly correlated with disease pathologies. Consistent with our findings, NR4A2 protein expression is reduced in post-mortem AD hippocampus at Braak stage VI (*33*).

**Figure 4.**
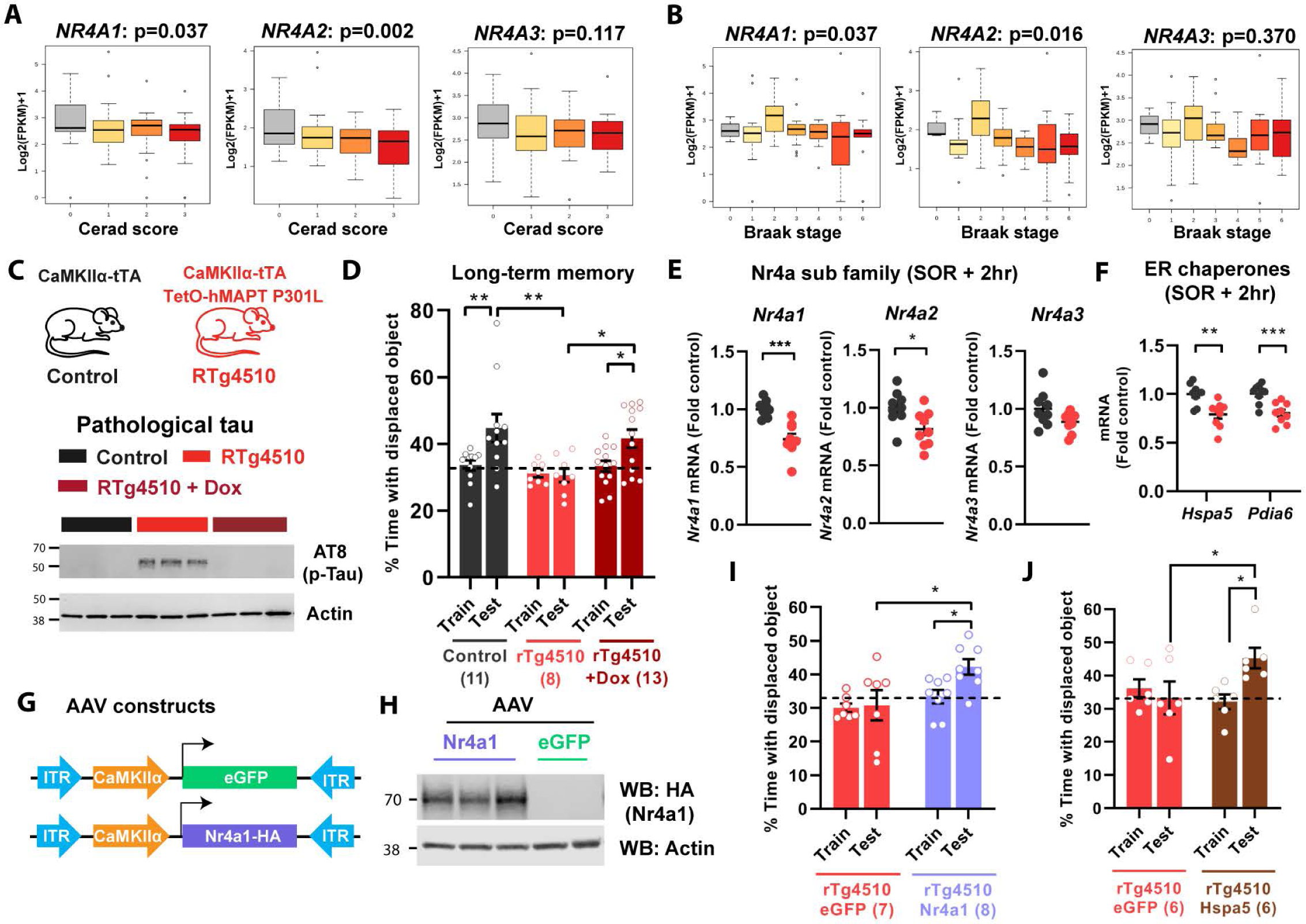
Restoration of Nr4a1 or ER chaperone function reverses memory deficits in a mouse model of ADRD. **(A-B)** Expression profiles of *NR4A1*, *NR4A2* and *NR4A3* in the hippocampus, from the Allen Brain Institute study of Aging, Dementia, and Traumatic Brain Injury, correlated with **(A)** Cerad scores, which reflect the density of neuritic plaques, and **(B)** Braak stages, which reflect the severity of neurofibrillary tangles. **(C)** Schematic depiction of control (CaMKIIα-tTA) or rTg4510 (CaMKIIα-tTA and TetO-hMAPT P301L) mice, and western blots of tau phosphorylation (AT8, phosphorylation at both Ser202 and Thr205) in the dorsal hippocampus in the presence or absence of doxycycline (Dox). **(D)** Long-term memory in rTg4510 and control mice at 4 mo of age, following training in SOR. Two-way ANOVA: significant main effect of genotype/treatment (F_(2, 29)_ =4.792, p=0.0159), significant main effect of sessions (F_(1, 29)_ =9.221, p=0.0050). Sidak’s multiple comparisons test: **p=0.0096 (control mice, Train versus Test), *p=0.0439 (rTg4510-Dox mice, Train versus Test), **p=0.0016 (rTg4510 mice, 24h test versus control mice, 24h test) and *p=0.0140 (rTg4510 mice, 24h test versus rTg4510 Dox, 24-hr test), control n=11 (3F), rTg4510 n=8 (4F) and rTg4510+Dox n=13 (7F). **(E-F)** rTg4510 and control mice (n=9 per group) were trained in SOR, and 2 hr later the dorsal hippocampus was collected and processed for RNA extraction and the analysis of gene expression. **(E)** *Nr4a* sub-family gene expression: Unpaired t-test: *Nr4a1*: t_(16)_ =4.878, ***p=0.0002; *Nr4a2*: t_(16)_ =2.621, *p=0.0185. **(F)** *Hspa5* and *Pdia6* gene expression: Unpaired t test: *Hspa5*: t_(16)_=3.692, **p=0.0020; *Pdia6*: t_(16)_=4.177, ***p=0.0007. **(G)** Schematic depiction of AAV constructs used to infuse into dorsal hippocampus of 3 mo-old rTg4510 mice. **(H)** Western blot showing expression of virus-transduced Nr4a1-HA in dorsal hippocampus 4 wk following infusion. **(I)** rTg4510 mice 3 mo of age were infused with AAV_9_CaMKIIα-Nr4a1-HA or control vector (AAV_9_-CaMKIIα-eGFP), and 4 wk later they were trained in SOR. Long-term memory was tested 24 hr after the training session. Two-way ANOVA: significant main effect of AAV-type infusion (F_(1, 13)_ =6.597, p=0.0234), Sidak’s multiple comparison tests: *p=0.0127 (AAV-Nr4a1, 24-hr test versus AAV-eGFP, 24-hr test) and *p=0.0489 (AAV-Nr4a1 Train versus AAV-Nr4a1 24-hr Test), AAV-Nr4A1: n=8 and AAV-eGFP: n=7. **(J)** rTg4510 mice 3.5 mo of age were infused with AAV_9_CaMKIIα-Hspa5-Tavi or control vector (AAV_9_-CaMKIIα-eGFP), and 2 wk later they were trained in SOR. Long-term memory was tested 24 hr after the training session. Two-way ANOVA: significant Session x AAV interaction F_(1, 10)_ =8.767, p=0.0143, Sidak’s multiple comparison tests: *p= 0.0419 (AAV-eGFP Test versus AAV-Hspa5 Test) and *p= 0.0419 (AAV-Hspa5 Train versus AAV-Hspa5 Test), AAV-eGFP: n=6 (3F) and AAV-Hspa5: n=6 (3F).

Cognitive impairment is a significant feature of ADRD, and we hypothesized that NR4A downregulation might be related to the compromise in cognitive abilities. To examine this in a mouse model of ADRD, we used the rTg4510 mouse, which overexpresses mutant human tau (tau P301L) exclusively in excitatory neurons (*35, 36*). These mice develop tangle-like inclusions (*35, 36*) and show pathological hyper-phosphorylation of tau proteins (AT8) in the dorsal hippocampus starting at 3-4 months age (**Fig. 4C**). These mice have deficits in spatial learning in the Morris water maze (MWM), contextual fear conditioning (*35–38*), and long-term spatial memory in the SOR task (**Fig. 4D**). Doxycycline treatment prevents these memory deficits (**Fig. 4D**), demonstrating that it is the expression of the mutant tau transgene and not the transgene insertion site (*39*) that drives the tauopathy-like phenotype. As in the case of the human post-mortem data, the expression of both *Nr4a1* and *Nr4a2* was downregulated in the dorsal hippocampus of rTg4510 mice after SOR training (**Fig. 4E**). These findings validate the appropriateness of utilizing this mouse line as a model for Nr4a dysregulation. Furthermore, we found that *Hspa5* and *Pdia6* were downregulated in the dorsal hippocampus of rTg4510 mice after SOR training (**Fig. 4F**). To determine the extent to which Nr4a transcription factors contribute to the memory impairment seen in rTg4510 mice, we overexpressed *Nr4a1* in the dorsal hippocampus of adult mice (**Fig. 4G-H**). This reversed the deficits in long-term spatial memory normally observed in rTg4510 mice (**Fig. 4I**). Lastly, to determine the role of Hspa5 chaperone in ADRD associated memory impairment, we overexpressed Hspa5 in the dorsal hippocampus of rTg4510 mice. Strikingly, Hspa5 overexpression ameliorated long-term memory deficits in rTg4510 mice (**Fig. 4J**). These findings link the function of the Nr4a family of transcription factors to the cognitive deficits associated with neurodegenerative disorders, and they suggest that targeting the Nr4a family and their downstream effector genes would be beneficial in the treatment of memory deficits associated with ADRD.

## Discussion

Here, we show that Nr4a transcription factors act during memory consolidation to drive the expression of genes encoding chaperones that are part of a multiprotein complex within the ER, thereby facilitating folding of the proteins into their functional conformations (*18*). This study provides functional evidence that these chaperones are involved in synaptic plasticity and long-term memory. The results demonstrating that long-term memory can be reversed in Nr4ADN mice by either application of the chemical chaperone PBA or overexpression of Hspa5 reveal that the protein folding machinery plays a critical role in memory consolidation. Our work in hippocampal neurons identifies the synaptic membrane protein GluN2A as a candidate target protein whose surface trafficking is regulated by Nr4a1-driven expression of Hspa5. Nr4a1 was previously shown to be involved in regulating dendritic spine density (*40*), consistent with the hypothesis that the target genes of this transcription factor impact synaptic structure and function. Our identification of ER chaperones as effector genes during memory consolidation provides a novel link between the induction of gene expression and protein synthesis, which are hallmarks of memory consolidation and the synaptic plasticity that leads to modification of neural circuits and behavioral alterations.

Our study extends these fundamental findings on the molecular mechanisms of memory, advancing our understanding of memory loss associated with neurodegenerative disorders by identifying changes in expression of the Nr4a family of transcription factors in both human AD brains and a mouse model of ADRD. Tau transgenic models and human tauopathy data exhibit widespread loss of heterochromatin (*41*), impaired chromatin remodeling and nuclear lamina formation (*42*). Levels of the lysine acetyl-transferase CBP are reduced in THY-Tau22 mutant mice (*3*) and in human AD patient samples (*43*). CBP and histone acetylation regulate the expression of Nr4a family genes, and consistent with our findings that overexpression of *Nr4a1* reverses memory deficits in tau mutant mice, both the pharmacological activation of CBP (*3, 44, 45*) and the inhibition of HDAC activity restore memory in several mouse models of ADRD (*46–48*). Although the exact mechanisms underlying the transcriptional alterations in neurodegenerative disorders remain to be identified, they represent attractive targets for the development of drugs to ameliorate cognitive deficits, which are a debilitating aspect of ADRDs. Our finding that overexpression of Nr4a1 or Hspa5 chaperone reverses memory loss in a tau-based model of ADRD supports this as a novel therapeutic approach.

The work described here identifies ER chaperone proteins as critical molecular regulators of memory storage. ER chaperones have been studied mainly for their roles in ER stress and the unfolded protein response; their role in memory consolidation is underexplored. Our work here links a subset of these ER chaperones, including Hspa5 and protein disulfide isomerases, to protein folding and trafficking within critical time windows during memory consolidation, laying the groundwork for future experiments to identify additional downstream targets of these ER chaperones, with promises of a more complete understanding of the fundamental molecular mechanisms of memory consolidation that go awry in neurodegenerative disorders.

## Acknowledgements

We thank the Iowa Institute of Human Genetics (IIHG) core for RNA seq library preparation and sequencing. We thank the Neural Circuits and Behavior Core in the Iowa Neuroscience Institute for use of their facilities. We thank Dr. Ron Merrill and Dr. Stefan Strack for their help with optimization of cell culture experiments and Dr. Lisa Lyons, Dr. Thomas Nickl-Jockschat, Dr. Joshua Weiner and Dr. Kevin Campbell for comments on the manuscript. We also thank Cindy Cosme, Samuel Dahlke, and Saaman Ghodsi for technical assistance.

## Author contributions

S.C. and T.A. conceived the study. S.C. and T.A. designed the experiments with input from J.M. and K.P.G. S.C. performed the behavioral tasks, stereotactic surgeries and molecular biology experiments. E.B. and Y.V. performed the bioinformatic analysis. U.M., J.D.L., A.L.Y. and E.N.W. performed biochemical experiments and analyzed behavioral results. U.M. performed cell culture experiments. M.S.S. performed electrophysiological experiments. S.C. and T.A. interpreted the results and wrote the article.

## Data and materials availability

The data that support the findings of this study are available within the article, its Extended Data files and Supplemental Files. All the uncropped western blots and raw data are also provided. The RNA seq data have been deposited in the NCBI Gene Expression Omnibus and are accessible through GEO Series accession number GSE167566. The code for analyses and figures related to RNA-seq data can be accessed through GitHub (https://github.com/ethanbahl/chatterjee2021_nr4a).

## Supplementary Materials

### Materials and Methods

#### Data reporting

No statistical methods were used to predetermine sample size.

#### Mouse lines

##### Nr4ADN mice

Adult Nr4ADN mice were 2-4 months old at the time of behavioral or biochemical experiments. They were maintained on a C57BL/6J background and harbor both the CaMKIIα-tTA and Tet-O-Nr4ADN transgenes(*1*). Incorporation of the CaMKIIα-tTA transgene into chromosome 12 causes a 508.12 Kb deletion that affects 5 genes: Vipr2, Wdr60, D430020J02Rik, Ncapg2 and Ptprn2. To account for any effects of the deleted genes on memory or gene expression(*2*), age-matched CaMKIIα-tTA expressing littermates were used as controls throughout the study.

##### rTg4510 mice

These mice were 3-4 months old at the time of behavioral or biochemical experiments. They were maintained on a C57BL/6J background and harbor two transgenes: CaMKIIα driven tTA, and TetO driven human tau P301L. Age-matched CaMKIIα-driven tTA expressing littermates were used as controls.

##### C57BL/6J mice

Adult male mice purchased from Jackson Laboratories were 2-4 months age during behavioral or biochemical experiments. All mice had free access to food and water; lights were maintained on 12 h: 12 h light/dark cycle. To suppress TetO driven expression of Nr4ADN or human tauP301L transgenes, Nr4ADN or rTg4510 mice were placed on a diet containing doxycycline (200 mg/kg, Bio-Serv) from weaning until behavioral experiments. All behavioral testing was performed during the light cycle between Zeitgeber time (ZT) 0-2. For all behavioral and biochemical experiments, mice were randomly assigned to groups, housed individually for seven days prior to experiments, and handled for 2 min per day for 5 days. All experiments were conducted according to US National Institutes of Health guidelines for animal care and use and were approved by the Institutional Animal Care and Use Committee of the University of Iowa, Iowa.

#### Drugs

Sodium phenyl butyrate (PBA, Sigma) and Sodium butyrate (NaBu, Sigma) was dissolved in saline. For electrophysiology experiments, the dose of PBA was chosen based on the range of IC_50_ values reported in the published literature(*3*). For behavioral experiments, mice were injected with PBA or NaBu *i.p.* at a dose of 200 mg/kg, immediately after SOR training. Control mice were injected with vehicle (0.9% sterile saline).

#### Adeno-associated virus (AAV) constructs

AAV_2.9_-CaMKIIα-eGFP, AAV_2.9_-CaMKIIα-Nr4A1-HA, AAV_2.9_-CaMKIIα-Hspa5-Tavi, AAV_2.2_-CaMKIIα-Nr4ADN, and AAV_2.2_-CaMKIIα-eGFP were purchased from VectorBuilder (VectorBuilder Inc).

#### Stereotactic surgeries

Mice were anaesthetized using isoflurane and kept on a warm heated pad throughout the stereotactic surgery procedure. Meloxicam was injected as analgesics(*4*). Viral infusion was performed using a 33G beveled needle (World Precision Instruments, WPI) attached to a 10 µl Nanofil syringe controlled by a microsyringe pump (UMP3; WPI). The coordinates for dorsal hippocampus were: anteroposterior, −1.9 mm, mediolateral, ±1.5 mm, and 1.5 mm below bregma. The needle was lowered to the site of injection over the course of 5 min and remained at the target for 1 min before injection was initiated (0.2µl per min). Each hippocampus was injected with approximately 1 µl of the relevant constructs. After injection was completed, the needle remained at the site for one additional minute and then slowly removed over a 5 min period. A small amount of bone wax (Lukens) was then used to close the drill holes and the incision was closed with sutures.

#### Spatial object recognition task

All animals were housed individually for 1 wk before behavioral experiments were initiated. Age-matched littermates were used. Mice were handled for 2 min per day for 5 consecutive days prior to the behavioral task. All spatial memory tasks were conducted between ZT0 to ZT2. Briefly, mice were habituated in the open field arena for 6 min during the habituation session, followed by three 6-min training sessions in the same open field containing three different glass objects. The intertrial interval was 3 min, during which the mice were returned to their homecage and the objects and arena were cleaned with 70% ethanol. An internal spatial cue (vertical black lines printed on a white paper 18 cm X 12 cm in size) was attached to one wall of the open field to allow the mice to locate each object relative to the spatial cue during free exploration of the arena. After either 1 hr or 24 hr following the training sessions, mice were brought back to the open field in which the location of one of the objects was displaced to a novel spatial location. Time spent exploring the displaced object (DO) and the non-displaced objects (NDO) during the 6-min test session was recorded. The exploration was hand-scored by an experimenter blinded to the genotype or treatment. Animals were assigned to the arenas randomly, without use of any randomization software. An object was scored as “explored” if it was sniffed or touched, or the face was in close proximity (within 1 cm) to the object, as described previously(*5*).

#### Electrophysiology

Nr4ADN male mice 2-3 months age were used. Mice were euthanized by cervical dislocation and the brain was quickly dissected into cold artificial cerebrospinal fluid (aCSF), which was continuously bubbled with carbogen (95% O_2_, 5% CO_2_). The isolation of hippocampi and preparation of acute hippocampal slices were performed as described(*6*). Transverse acute hippocampal slices of 400-µm thickness were prepared from both hippocampi using a manual McIlwain slicer (Stoelting). The slices were quickly transferred onto a net insert in an interface recording chamber (Fine Science Tools, Foster City, CA) and left to equilibrate to a humidified carbogen atmosphere at 28°C for at least 2-3 hr before recordings were initiated. The slices were perfused at 1 mL/min with oxygenated aCSF throughout the experiments. The aCSF used for both the dissection and recordings was composed of 124 mM NaCl, 4.4 mM KCl, 1 mM NaH_2_PO_4_, 2.5 mM CaCl_2_.2H_2_O, 1.3 mM MgSO_4_.7H_2_O, 26.2 mM NaHCO3 and 10 mM D-glucose; pH ∼7.4 when equilibrated with carbogen. Field excitatory post-synaptic potentials (fEPSPs) were recorded in the CA1 stratum radiatum by stimulating Schaffer collaterals with a monopolar, lacquer-coated stainless-steel electrode ((∼5 MΩ resistance, A-M Systems, # 571000) and recording with an aCSF-filled glass microelectrode (2–5 MΩ resistance). In all experiments, test stimulation was a biphasic, constant-current pulse (100 µs duration) delivered every min at a stimulation intensity that evoked ∼40% of the maximal fEPSP amplitude, as determined by an input-output curve (stimulation intensity vs fEPSP amplitude). Also, a stable baseline was recorded for at least 20 min before LTP was induced or drug was applied. LTP was induced by a spaced 4-train stimulation protocol consisting of four 100 Hz, 1-sec trains delivered at 5-min intervals, at the test stimulus intensity. For each experiment, PBA solution (2 mM) was prepared fresh by dissolving in aCSF, and it was applied to the bath and protected from light. The solution was recirculated after 30 min of initial application. In the electrophysiological data presented, ‘n’ represents the number of slices. Data were acquired using Clampex 10 and Axon Digidata 1440 digitizer (Molecular Devices, Union City, CA) at 20 kHz and were low-pass filtered at 2 kHz with a four-pole Bessel filter. Data analysis was performed using the Clampfit 10 software (Molecular Devices, Union City, CA). Data were plotted and statistical analyses were performed using the GraphPad Prism 8 software. For each slice, the fEPSP slopes were normalized against the average slope over the 20-min baseline (pre-drug baseline in the PBA-treated slices). Data are presented as mean ± SEM. LTP persistence was assessed by comparing the final 20-min recordings for the vehicle and treatment groups using a two-way repeated measures ANOVA. The mean fEPSP slope of the final 20-min recordings for each group was compared using two-tailed, unpaired t-test. Statistical significance was set at p<0.05.

#### Isolation of whole-cell extracts and synaptosomal fractions

For whole-cell lysate preparation, flash frozen dorsal hippocampal tissue was homogenized mechanically in 300 µl of ice-cold RIPA buffer (Sigma) supplemented with 0.2% Triton X-100 (Sigma), and Protease and Phosphatase Inhibitor Cocktail (1:100, Thermo Scientific). The lysate was kept on ice for 30 mins, following which they were centrifuged at 10,000 x g for 15 min at 4^0^C. The pellet was discarded, and the supernatant (whole cell lysate) was collected for Western blot analysis. For synaptosomal extraction, Hippocampal tissue was mechanically homogenized in Syn-PER Reagent (Thermo Fisher Scientific) containing Halt Protease and Phosphatase Inhibitor Cocktail (1:100, Thermo Scientific). The homogenate was centrifuged at 1200 x g for 10 min at 4°C, after which the pellet (nuclear fraction) was discarded and the supernatant was centrifuged again at 15,000 x g for 25 min at 4°C. This pellet (synaptosomal fraction) was resuspended in RIPA buffer (Sigma) containing Halt Protease and Phosphatase Inhibitor Cocktail (1:100) and 0.2% Triton X-100 (Sigma).

#### Western blot analysis

Protein extracts were transferred to polyvinylidene difluoride membranes as previously described(*7*). Membranes were blocked with Odyssey® Blocking Buffer in TBS (LI-COR) and incubated overnight at 4°C with the following primary antibodies: Hspa5 (1:2000, Proteintech 11587-1-AP), Pdia6 (1:2000, Abcam ab11432), Biotin Ligase Epitope Tag (for Tavi-tag detection, 1:1000, Abcam ab106159), PSD95 (1:5000, ThermoFisher Scientific, 6G6-1C9), HA (1:2000, Millipore Sigma), phospho-tau AT8 (1:5000, Biolegend, 806503), Actin (1:10,000, ThermoFisher Scientific), ATF4 (1:500, Thermo Fisher Scientific, PA5-27576), ATF6 (1:500, Thermo Fisher Scientific, PA5-114886), phospho-IRE1alpha (1:500, Thermo Fisher Scientific, PA5-85738). Membranes were washed and incubated with appropriate IRDye IgG secondary antibodies, including anti-rabbit IRDye 800LT (1:5,000, LI-COR) and anti-mouse IRDye 680CW (LI-COR). Images were acquired using the Odyssey Infrared Imaging System (LI-COR). Quantification of western blot bands was performed using Image Studio Lite ver5.2 (LI-COR).

#### RNA extraction, cDNA synthesis and quantitative real-time reverse transcription (RT)-PCR

Dorsal hippocampi were dissected and immediately stored at -80°C in RNAlater solution (Ambion) for later isolation of total RNA. For RNA extraction, Qiazol (Qiagen) was added to the hippocampal tissues and they were homogenized using stainless steel beads (Qiagen). Chloroform was then added to the homogenates and the samples were centrifuged at 12,000 x g at RT for 15 min. RNA was precipitated from the aqueous phase using ethanol and then cleaned using the RNeasy kit (Qiagen). RNA was eluted in nuclease-free water and treated with DNase (Qiagen) at RT for 25 min. The cleaned RNA was precipitated in ethanol, sodium acetate (pH 5.2) and glycogen overnight at -20°C. RNA samples were centrifuged at top speed at RT for 20 min. The precipitates were further washed with 70% ethanol and centrifuged at top speed for 5 min. The RNA precipitates were dried and resuspended in nuclease free water, and concentrations were estimated using a Nanodrop (Thermo Fisher Scientific). cDNAs were prepared from 1 µg RNA using the SuperScript™ IV First-Strand Synthesis System (Ambion). Real-time RT-PCR reactions were performed in a 384-well optical reaction plate with optical adhesive covers (Life Technologies). Each reaction was composed of 2.25μl cDNA (2 ng/ul), 2.5μl Fast SYBR™ Green Master Mix (Thermo Fisher Scientific), and 0.25μl of primer mix (IDT). A minimum of three technical replicates per reaction was performed on the QuantStudio 7 Flex Real-Time PCR system (Applied Biosystems, Life Technologies). Data was normalized to housekeeping genes (*Tubulin*, *Pgk1* and *B2m*) and 2^(-ΔΔCt)^ method was used for gene expression analysis.

#### RNA library preparation, sequencing and analysis

RNA libraries were prepared at the Iowa Institute of Human Genetics (IIHG), Genomics Division, using the Illumina TruSeq Stranded Total RNA with Ribo-Zero gold sample preparation kit (Illumina, Inc., San Diego, CA). KAPA Illumina Library Quantification Kit (KAPA Biosystems, Wilmington, MA) was used to measure library concentrations. Pooled libraries were sequenced on Illumina HiSeq4000 sequencers with 150-bp Paired-End chemistry (Illumina) at the IIHG core. RNA-sequencing data was processed with the bcbio-nextgen pipeline (https://github.com/bcbio/bcbio-nextgen, version 1.1.4). The pipeline uses STAR(*8*) to align reads to the mm10 genome build (GENCODE release M10, Ensembl 89 annotation) and quantifies expression at the gene level with featureCounts(*9*). All further analyses were performed using R(*10*). For gene level count data, the R package EDASeq(*11*) was used to adjust for GC content effects (full quantile normalization) and account for sequencing depth (upper quartile normalization) (Extended Data Fig 10 and 12). Latent sources of variation in expression levels were assessed and accounted for using RUVSeq (RUVr mode using all features)(*12*) (Extended Data Fig 11 and 13). Appropriate choice of the RUVSeq parameter *k* was guided through inspection of *P* value distributions, relative log expression (RLE) plots, and principal component analysis (PCA) plots. Specifically, the smallest value *k* was chosen where the *P* value distribution showed an expected peak below 0.05, RLE plots were evenly distributed and zero-centered, and PCA plots demonstrated replicate sample clustering in the first three principal components(*13*). Differential expression analysis was conducted using the edgeR quasi-likelihood pipeline(*14–16*). Codes to reproduce the RNA-sequencing analysis are available at https://github.com/ethanbahl/chatterjee2021_nr4a.

#### GO and pathway enrichment analyses of DEGs

The ClueGO(*17*) and CluePedia plug-ins of the Cytoscape 3.7.5 software(*18*) were used in “Functional analysis” mode, using the default parameters for analyzing gene ontology, molecular function and KEGG pathways in networks for DEGs. The names of significant DEG were pasted into the “Load Marker List” of ClueGO, and the organism “Mus Musculus [10090]” was selected.

#### Construction of protein-protein interaction (PPI) networks

The protein-protein interactive network was constructed using STRING(*19*) (version 11.0), which uses the STRING database (http://string-db.org/)(20). The PPI network was constructed to identify the interactions between proteins encoded by down-regulated DEGs based on experimental data. The DEG names were pasted into “STRING protein query”. Active interaction sources, including text-mining, experiments, databases, co-expression, neighborhood, gene fusion and co-occurrence were applied and highest interaction score confidence (0.900) was selected to construct the PPI networks. Full network was constructed, where the edges indicate both functional and physical protein associations.

#### ChIP-seq analysis

Using the SRA toolkit fastq-dump, raw ChIP-seq data from Liu et al(*21*) (GEO accession code GSE96969) was downloaded, including Nr4a1-HA (SRR6788331), IgG-control (SRR6788333), and input DNA (SRR6788332). The raw reads were trimmed using Trimmomatic version 0.36 with the parameters ILLUMINACLIP:2:30:15 LEADING:30 TRAILING:30 MINLEN:23, and quality inspection was conducted using FastQC version 0.11.5. The trimmed reads from all data sets were aligned using BWA version 0.7.15 with the algorithm mem and default parameters. Signal density files in BedGraph format were generated using BEDTools genomecov version 2.26.0 with default parameters and then converted in uniform 10 nucleotide-bin WIG files for further normalization steps. Peak calling was done using MACS2 version 2.1.2 with a q-value cutoff of 0.01 and default parameters. Each signal density file corresponding to a data set was scaled such that the total sum of the signal over the mouse mm10 genome was equivalent to 1M reads of 100 nucleotides. The signal of the input data set was then subtracted from its corresponding IP data set to generate the “sclWT-ctrl” files. The normalized WIG files were then encoded in bigWig format using the Kent utilities and visualized using the Integrative Genomics Viewer with the mouse mm10 reference genome.

#### Primary neuronal cultures and AAV-transduction

Hippocampi from mice (C57BL/6J) were used to generate primary neuronal cultures as previously described(*22*). Briefly, hippocampi from postnatal P0/P1 pups were dissected, trypsinized (0.25%), and triturated using a fire-polished glass Pasteur pipette to prepare a single-cell suspension. Cells were then plated on poly-L-lysine (1mg/ml) coated 4-well glass-bottom dishes (Cellvis) at optimal density (150-200 cells/mm^2^) and maintained in Neurobasal medium (Gibco) containing B27 Supplement (Gibco), in an incubator with 5% CO_2_ and at 37°C. At Days *in vitro* (DIV) 16-17, neurons were transduced with AAV_2.2_-CaMKIIα-Nr4ADN, AAV_2.9_-CamKIIa-Hspa5-Tavi, and and AAV_2.2_-CaMKIIα-eGFP (titer of concentrated viral stock was 1-2 × 10^13^TU/ml) in a 1:1000 dilution of Neurobasal Medium containing B27 Supplement. At 8-10 hrs following transduction, half of the existing medium was replenished with fresh Neurobasal medium containing B27 Supplement. Cultures were typically maintained until DIV 23-25 before experiments commenced.

#### KCl stimulation and surface labelling of GluN2A

At DIV 23-25, neurons were incubated in low KCl-HBS (290 mOSm) (110 mM NaCl, 5.4 mM KCl, 1.8 mM CaCl_2_, 0.8 mM MgCl_2_, 10 mM D-glucose, 10 mM HEPES-NaOH pH 7.4) for 60 mins. Thereafter, neurons were stimulated for another 60 mins with high KCl-HBS (same as low KCl-HBS, except for 55 mM NaCl and 60 mM KCl). The high KCl-HBS was washed off, and live neurons were then immunolabelled with N-terminal NMDAR2A antibody (1:25, Thermo Fisher) in low KCl-HBS for 30 mins to exclusively stain the surface GluN2A receptors. Following the antibody incubation, the cells were washed twice with phosphate buffered saline containing Mg^2+^and Ca^2+^ (PBS-MC; 137mM NaCl, 2.7 mM KCl, 10 mM Na_2_HPO_4_, 2mM KH_2_PO_4_, 1 mM MgCl_2_ and 0.1 mM CaCl_2_). Cells were then fixed in PBS-MC containing 2% paraformaldehyde and 2% sucrose for 15 minutes at 37°C, washed thrice in PBS-MC at room temperature and blocked with PBS-MC containing 2% BSA for 60 minutes at room temperature. Cells were incubated with Alexa-647 conjugated goat-anti-rabbit secondary antibody (1:200, Invitrogen) at room temperature for 90 minutes in blocking solution. Neurons were then permeabilized with PBS-MC containing 0.1% Triton-X-100 at room temperature for 5 minutes, incubated with blocking solution for 60 minutes, and thereafter with MAP2 antibody (1:1000, Sigma) for 8-10 hours at 4°C. For PSD95 immunocytochemistry, permeabilized neurons were incubated with PSD95 antibody (1:4000, Enzo Life Sciences) for 8-10 hours at 4°C. Cells were washed thrice in PBS-MC and incubated with Alexa-546 conjugated goat-anti-mouse secondary antibody (1:200, Invitrogen) at room temperature for 60 minutes. Finally, cells were washed three times with PBS-MC at room temperature and preserved in PBS-MC for future imaging.

#### Confocal imaging and image analysis

Cultured neurons after completion of the experiments were imaged using Olympus FV3000 confocal microscope with a 100X NA = 1.45 oil immersion objective at 1024 × 1024-pixel resolution. High magnification images were captured using 3X optical zoom. All images (8 bit) were acquired with identical settings for laser power, detector gain and pinhole diameter for each experiment and between experiments. Images from the different channels were stacked and projected at maximum intensity using ImageJ (NIH). Mean Fluorescence Intensity (MFI) of surface GluN2A and the colocalization between surface GluN2A and PSD95 punctas was assessed using plugins in ImageJ.

#### Analysis of human Alzheimer’s disease

Using RNA-sequencing data from the “Aging, Dementia and TBI Study”(*23*) provided by the Allen Institute for Brain Science, we fit linear models between the RIN-corrected and log2-transfsormed hippocampal expression levels of NR4A family members and the individual’s Cerad score, a semiquantitative estimate of neuritic plaque density(*23*).

#### Statistics

Behavioral and biochemical data were analyzed using paired or unpaired two-tailed t-tests and either one-way or two-way ANOVAs (in some cases with repeated measures as the within subject variable). Sidak’s tests were used for post-hoc analyses where needed. Differences were considered statistically significant when p<0.05. As indicated for each figure panel, all data are plotted in either bar graphs, in which symbols represent each data point, or in dot plots, where each symbol represents an individual data point. Graphs were plotted as mean ± SEM.

---

**Fig. S1.**
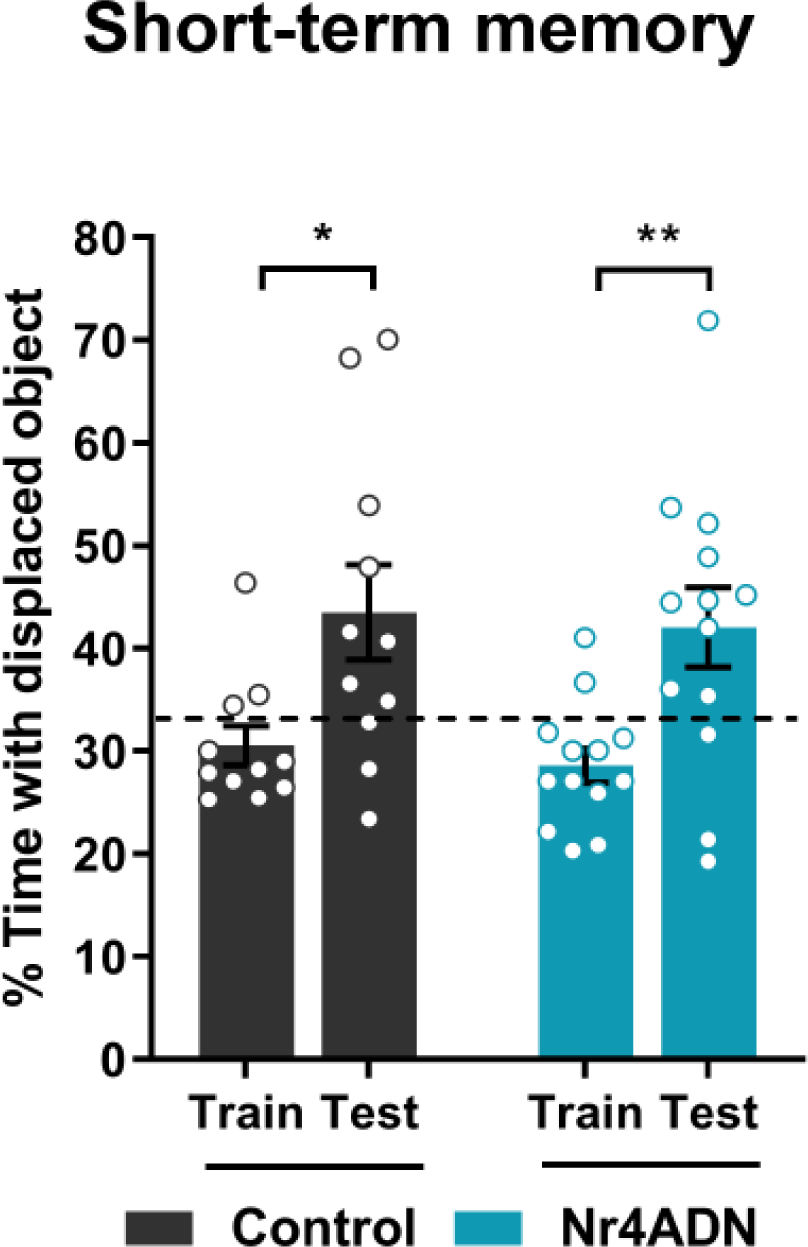
Nr4ADN mice show intact short-term memory. Both Nr4ADN and control mice showed significant preference for DO during the 1h short-term memory test. Two-way ANOVA: Significant main effect of session (F (1, 22) = 21.49, p<0.0001), Sidak’s multiple comparison tests: *p=0.0107 (control mice, train vs 24h test, n=11(5F)) and **p=0.0041 (Nr4ADN mice, train Vs 24h test, n=13 (7F)).

**Fig. S2.**
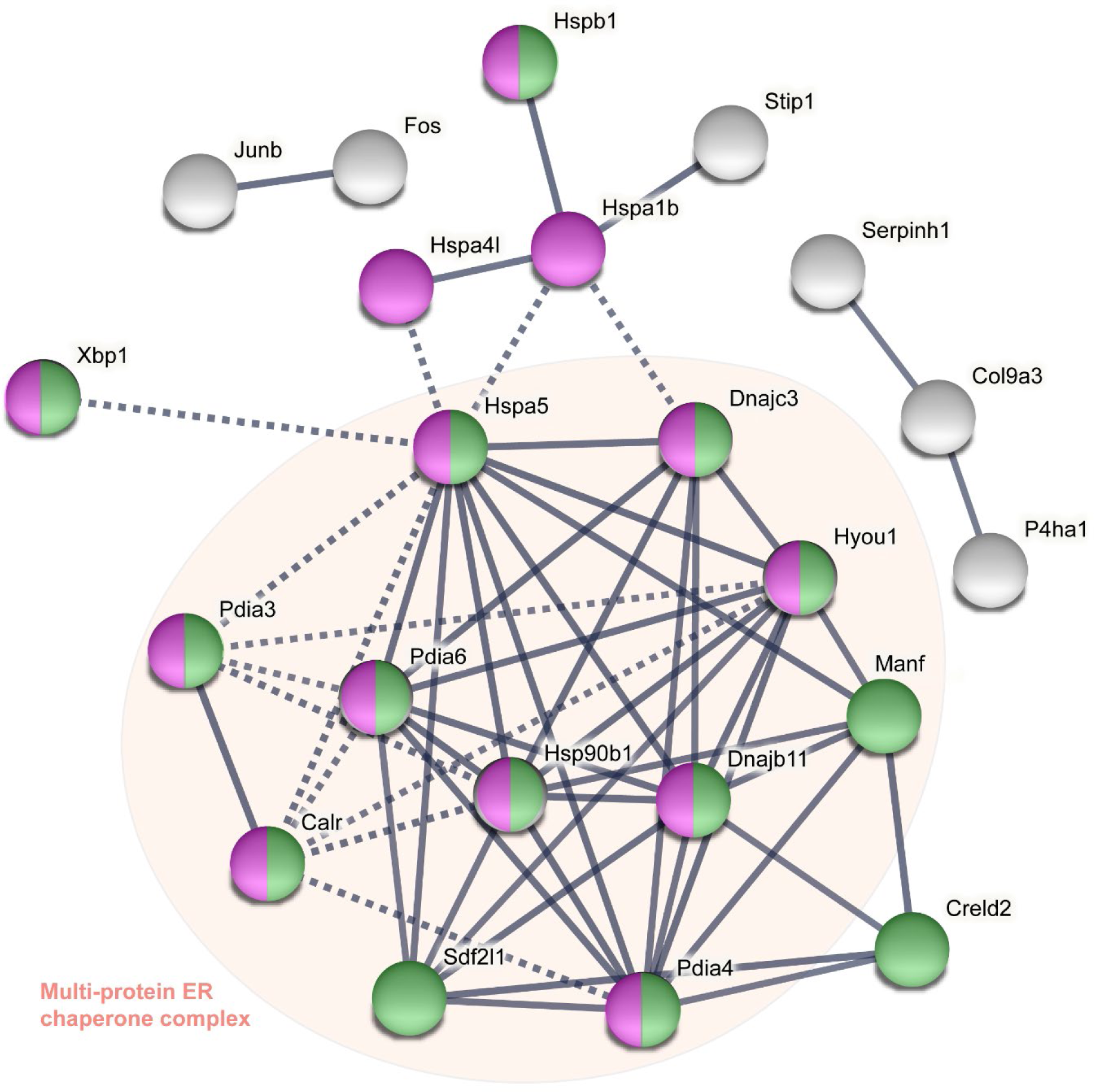
Nr4A transcription factors regulate the expression of genes encoding chaperone proteins that form a multiprotein ER chaperone complex. Depiction of multi-protein ER chaperone complex identified by STRING (Search Tool for the Retrieval of Interacting Genes/Proteins) analysis. Functional protein association networks were inferred from downregulated genes in Nr4ADN mice after learning. This complex consists of Hspa5, Pdia6 and other ER chaperones, and enables the proper folding and trafficking of nascent proteins. Green: proteins associated with ER chaperone complex (local network cluster in STRING, FDR=1.00e-26), Purple: proteins associated with protein processing in ER (KEGG pathway, FDR=3.29e-18). Edges represent confidence of protein-protein associations. Proteins inside the red shape are part of multiprotein ER chaperone complex which facilitates folding of nascent proteins.

**Fig. S3.**
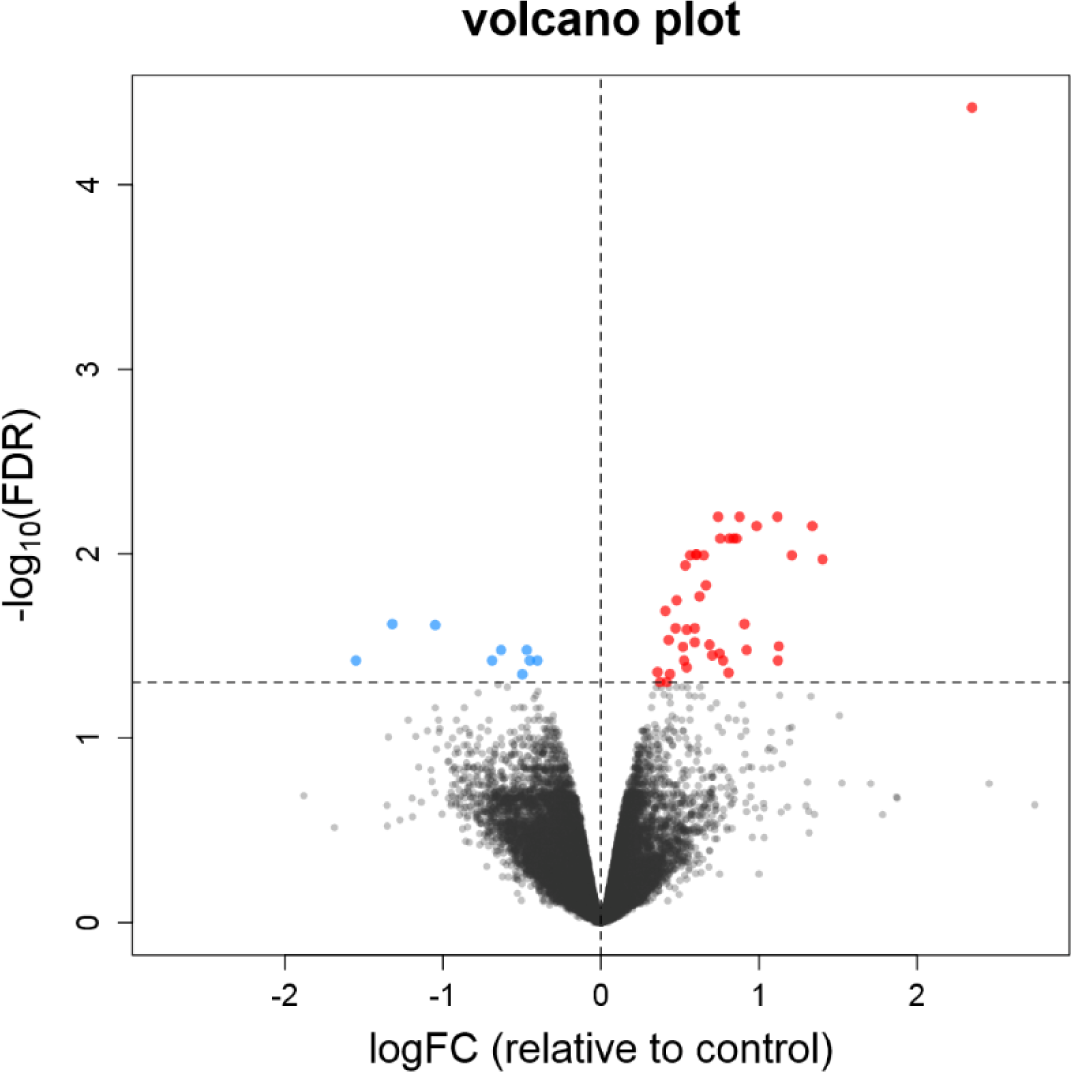
RNA-seq analysis for control mice. Volcano plot comparing genes that are differentially expressed in the dorsal hippocampus between control mice (tTA^+^ DN^-^ n=2, tTA^-^ DN^-^ n=2) 2 hr after SOR training and homecage controls (tTA^+^ DN^-^ n=2, tTA^-^ DN^-^ n=2). SOR learning led to the upregulation of 42 genes and the downregulation of 9 genes.

**Fig. S4.**
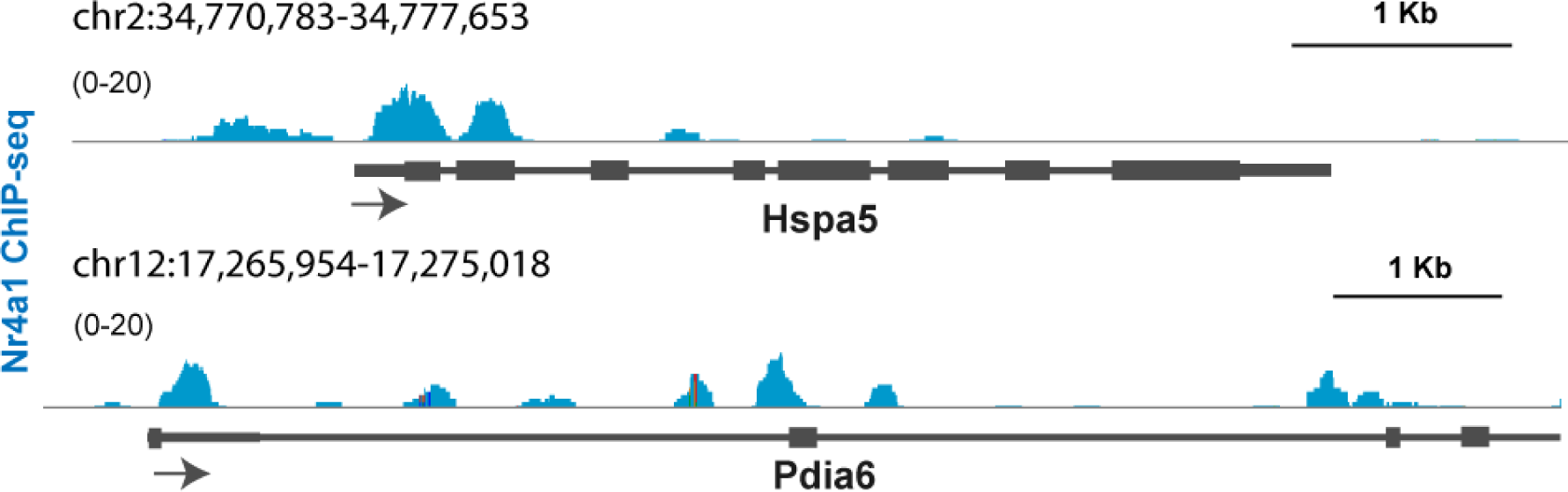
Nr4a1 is enriched on Hspa5 and Pdia6 promoter. Genome browser track view of ChIP-seq data for Nr4a1 peak at the promoters of *Hspa5* and *Pdia6* (Liu, X., et al. 2019).

**Fig. S5.**
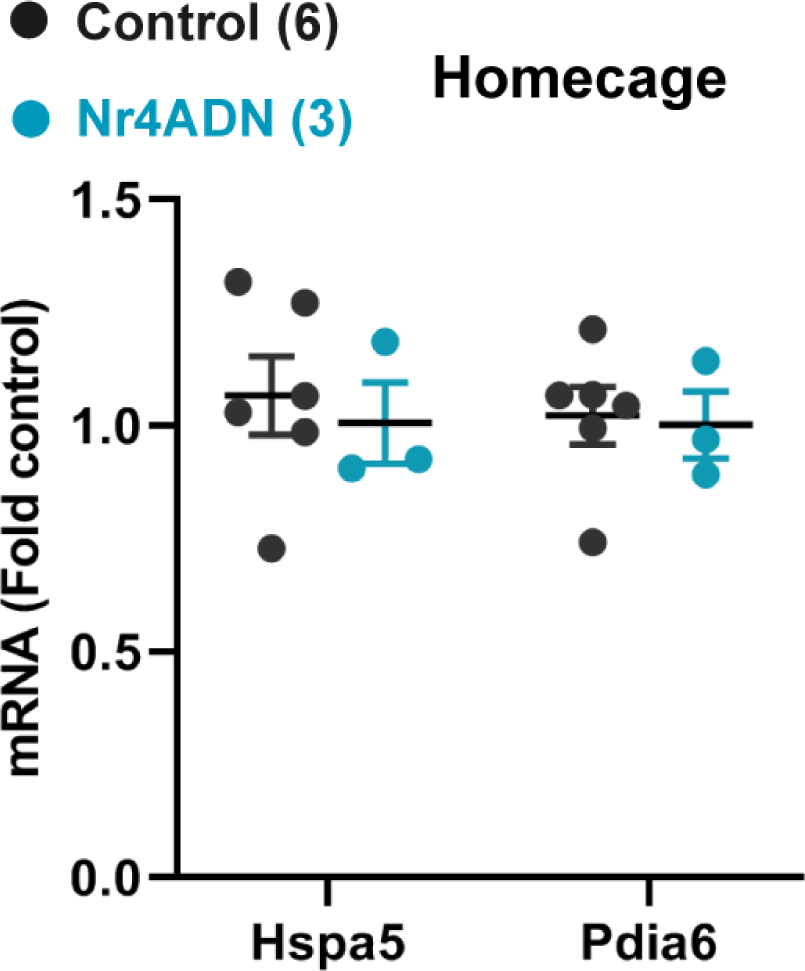
ER chaperones are not differentially expressed in homecage Nr4ADN mice. Levels of mRNAs encoding *Hspa5* and *Pida6* in 2 mo-old control and Nr4ADN in home cage. The dorsal hippocampus was isolated when the mice reached 2 mo of age and total RNA was extracted and analyzed by qPCR. Both the *Hspa5* and *Pdia6* mRNAs were expressed at similar levels in Nr4ADN and control mice. Unpaired t-test: t_(7)_=0.4304, p=0.6798 (*Hspa5*) and t_(7)_=0.1937, p= 0.8519 (*Pdia6*).

**Fig. S6.**
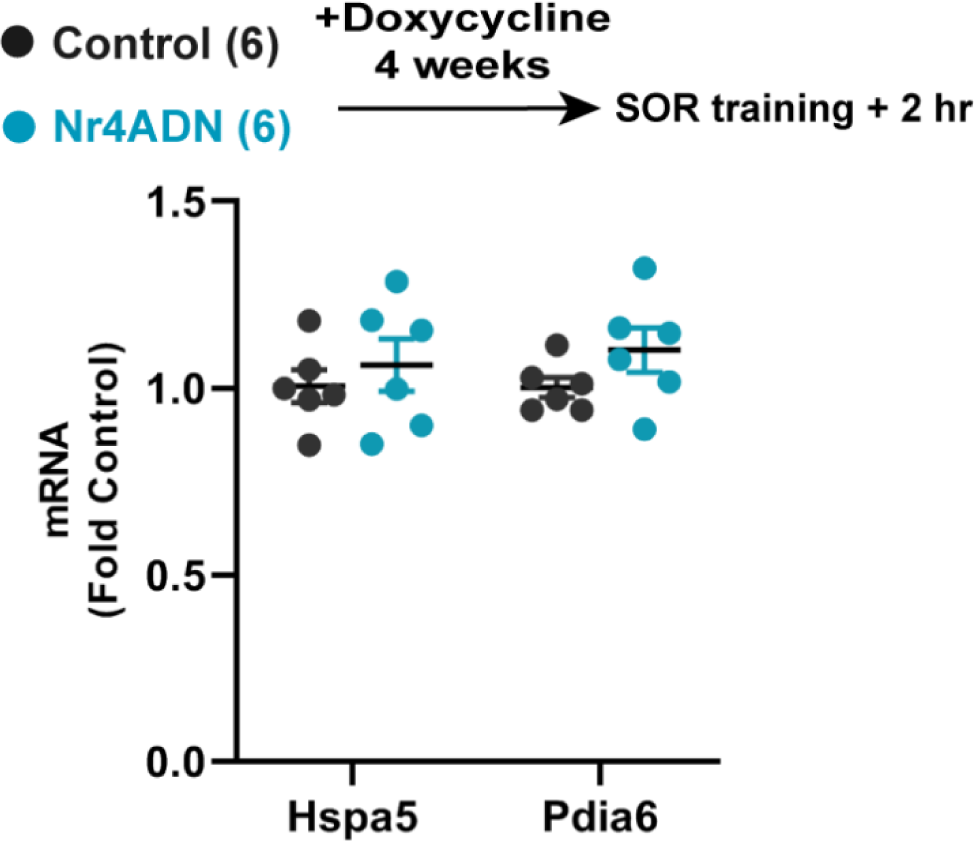
Blocking Nr4ADN transgene expression abolishes downregulation of ER chaperone genes. Levels of mRNAs encoding Hspa5 and Pida6 at 2 mo of age, after SOR training. Mice were placed on a diet containing doxycycline from weaning until 2 mo of age, to suppress transgene expression in the Nr4ADN mice. The mice were then trained in SOR and the dorsal hippocampus was removed 2 hr after training was completed. Total RNA was isolated, and qPCR was performed. *Hspa5* and *Pdia6* mRNA levels were equivalent to those in control littermates fed the same diet. Unpaired t-test: t_(10)_=0.6846, p=0.3338 (*Hspa5*), t_(10)_=1.536, p=0.1089 (*Pdia6*).

**Fig. S7.**
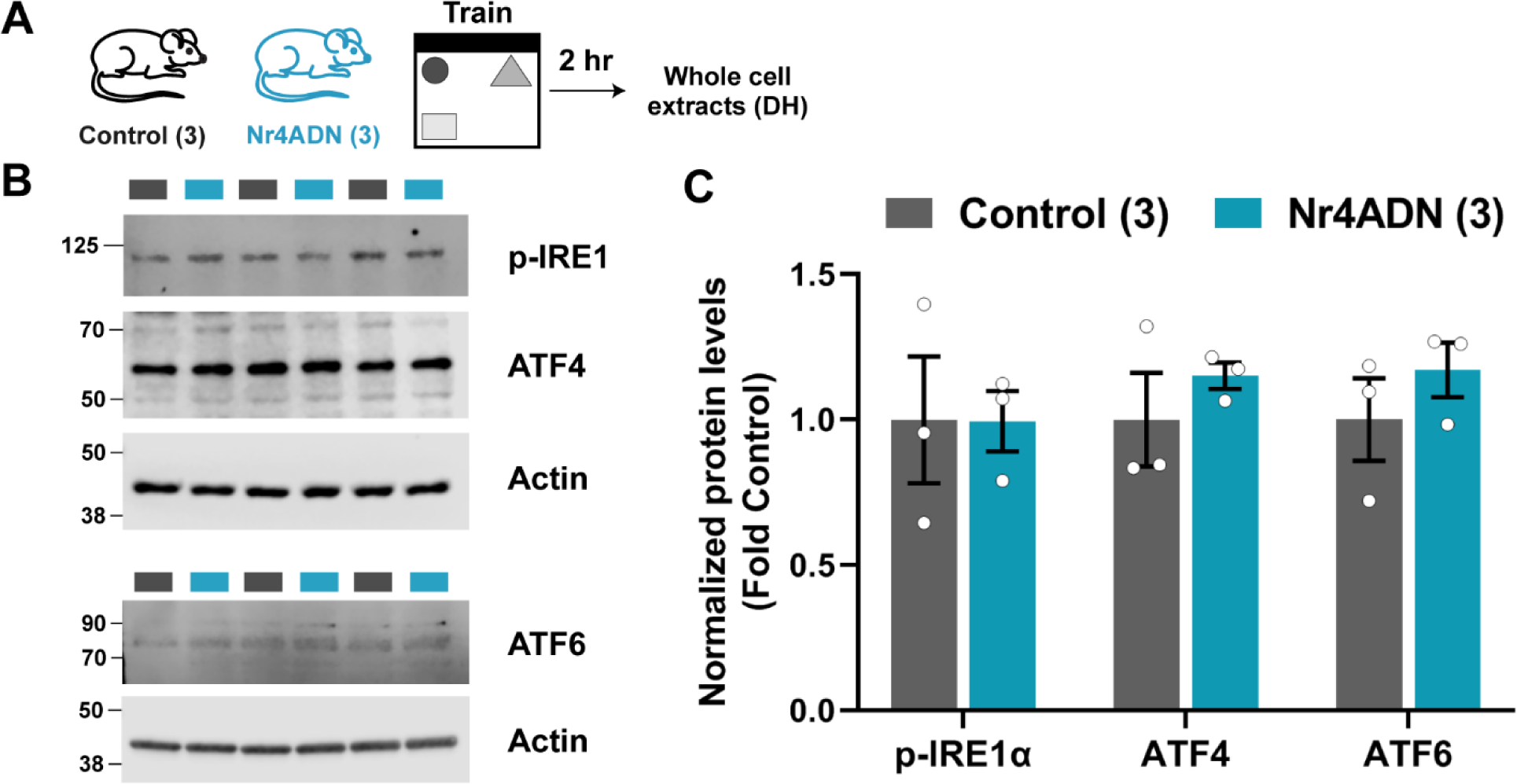
Nr4ADN mice do not show any evidence of elevated ER stress response following SOR learning. **(A)** Nr4ADN and control mice were trained in SOR and 2 hr after training, the dorsal hippocampus was extracted for analyses of ER stress markers. **(B)** p-IRE1, ATF4 and ATF6 levels were measured using Western blot analysis from whole cell extracts. **(C)** Quantification of p-IRE1, ATF4 and ATF6 expression levels after normalization to actin expression levels.

**Fig. S8.**
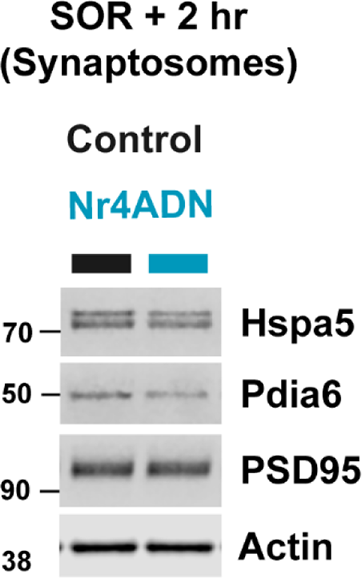
Nr4a regulates Hspa5 and Pdia6 protein levels. Western blot showing expression of Hspa5 and Pdia6 proteins from synaptosomes obtained from dorsal hippocampus of Nr4ADN and control mice 2 hr after SOR training.

**Fig. S9.**
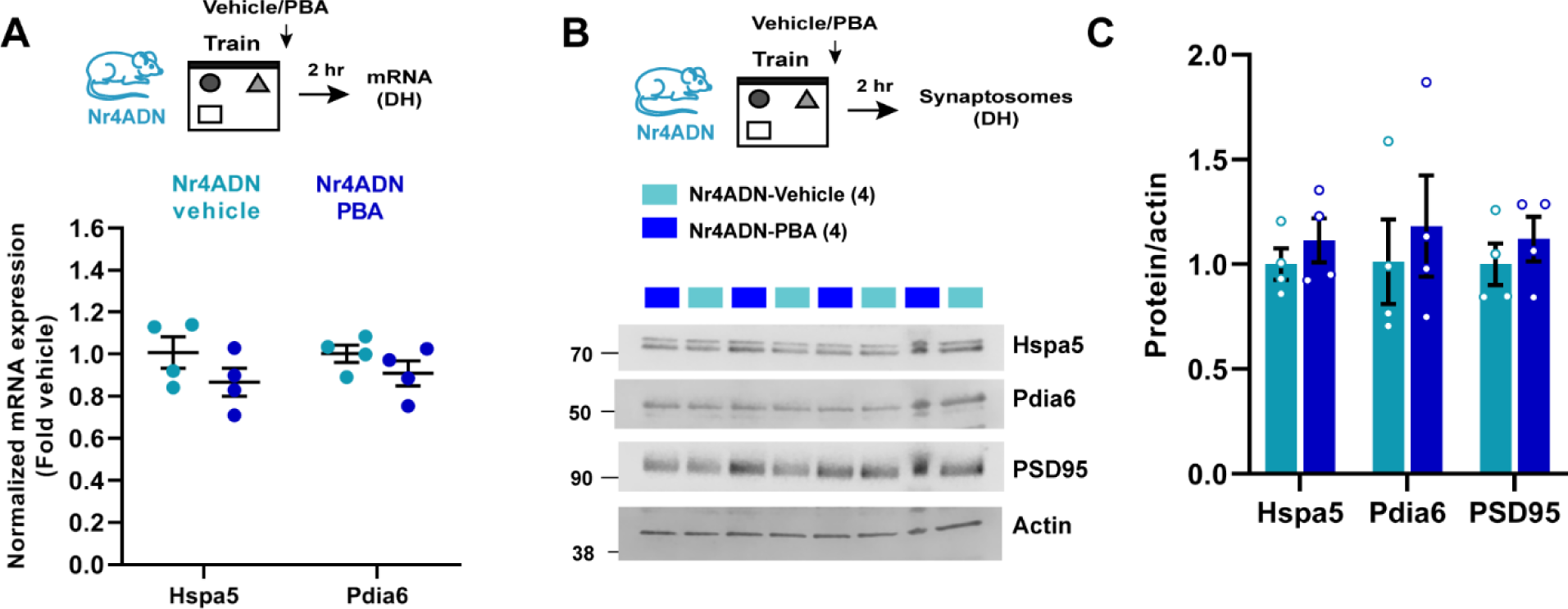
PBA treatment does not alter the expression of ER chaperones Hspa5 and Pdia6 in Nr4ADN mice. **(A-C)** Nr4ADN mice were trained in SOR and injected with either PBA (200 mg/kg) or vehicle (saline) immediately after the training session. 2 hr after training, the dorsal hippocampi were extracted for analyses of gene expression and synaptosomal proteins. **(A)** Expression of the mRNAs encoding the ER chaperones *Hspa5* and *Pdia6* was unaltered between PBA- and saline-injected Nr4ADN mice. Unpaired t-test: t_(6)_=1.402, p=0.8521 (*Hspa5*) and t_(6)_=1.290, p=0.5671 (*Pdia6*). **(B)** Western blot of synaptosomal extracts. **(C)** Quantification of data from **B** showing that expression levels of Hspa5 and Pdia6 proteins are similar in PBA- and saline-injected Nr4ADN mice. Unpaired t-test: t_(6)_=0.8800, p=0.5878 (Hspa5) and t_(6)_=0.5387, p=0.7688.

**Fig. S10.**
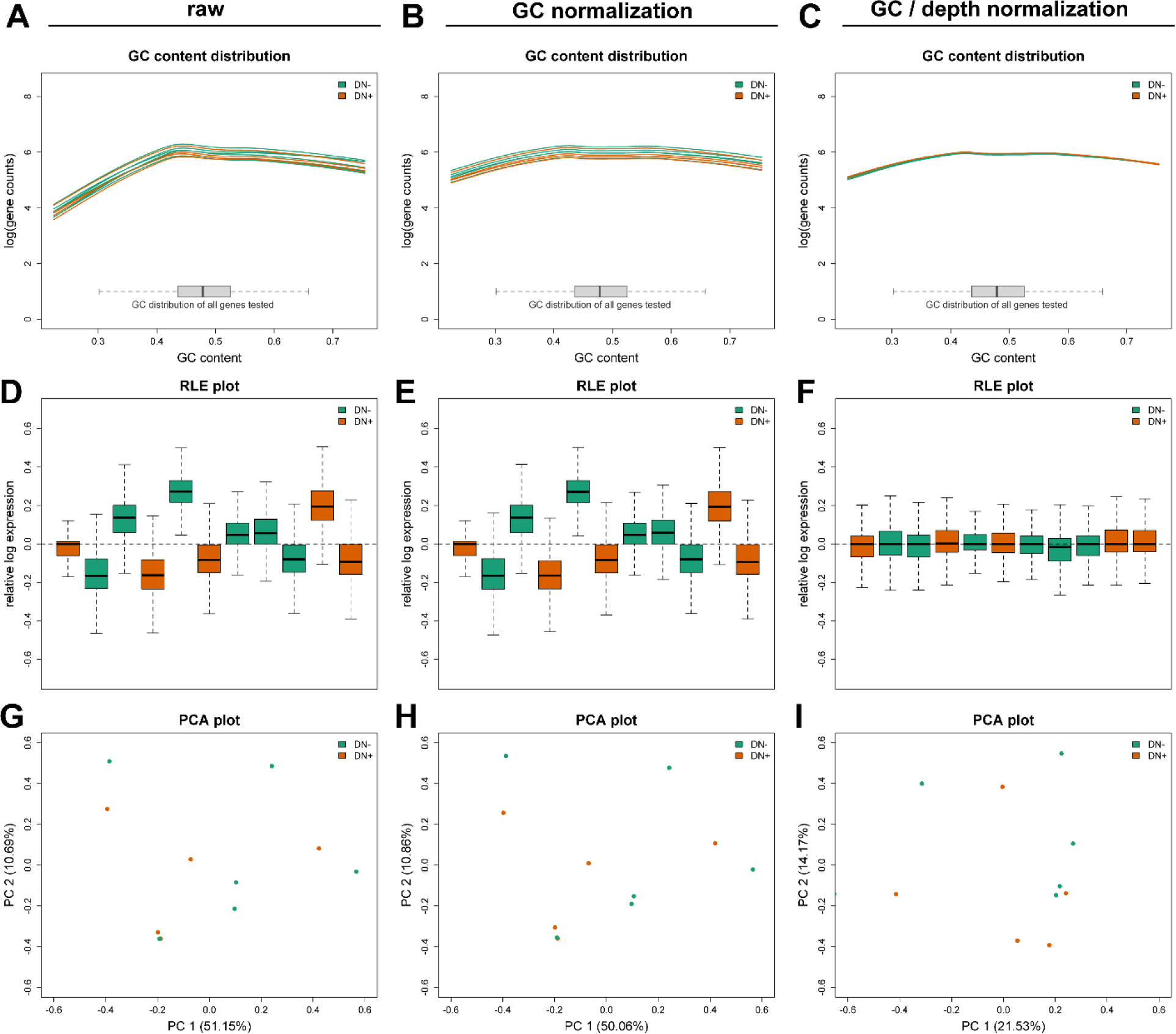
Normalization of differences in the distribution of GC content and sequencing depth using Exploratory Data Analysis and Normalization for RNA-Seq (EDASeq) for comparisons of RNAseq data for Nr4ADN and control mice after learning. **(A-C)** GC content distributions before normalization (**A**), after full quantile GC content normalization (**B**), followed by upper quartile sequencing depth normalization (**C**). **(D-F)** Relative log expression (RLE) plots before normalization (**D**), after full quantile GC content normalization (**E**), followed by upper quartile sequencing depth normalization (**F**). **(G-I)** Principal component analysis (PCA) plots before normalization (**G**), after full quantile GC content normalization (**H**), followed by upper quartile sequencing depth normalization (**I**). DN- (Control: tTA+, Nr4ADN-) and DN+ (Nr4ADN: tTA+, Nr4ADN+).

**Fig. S11.**
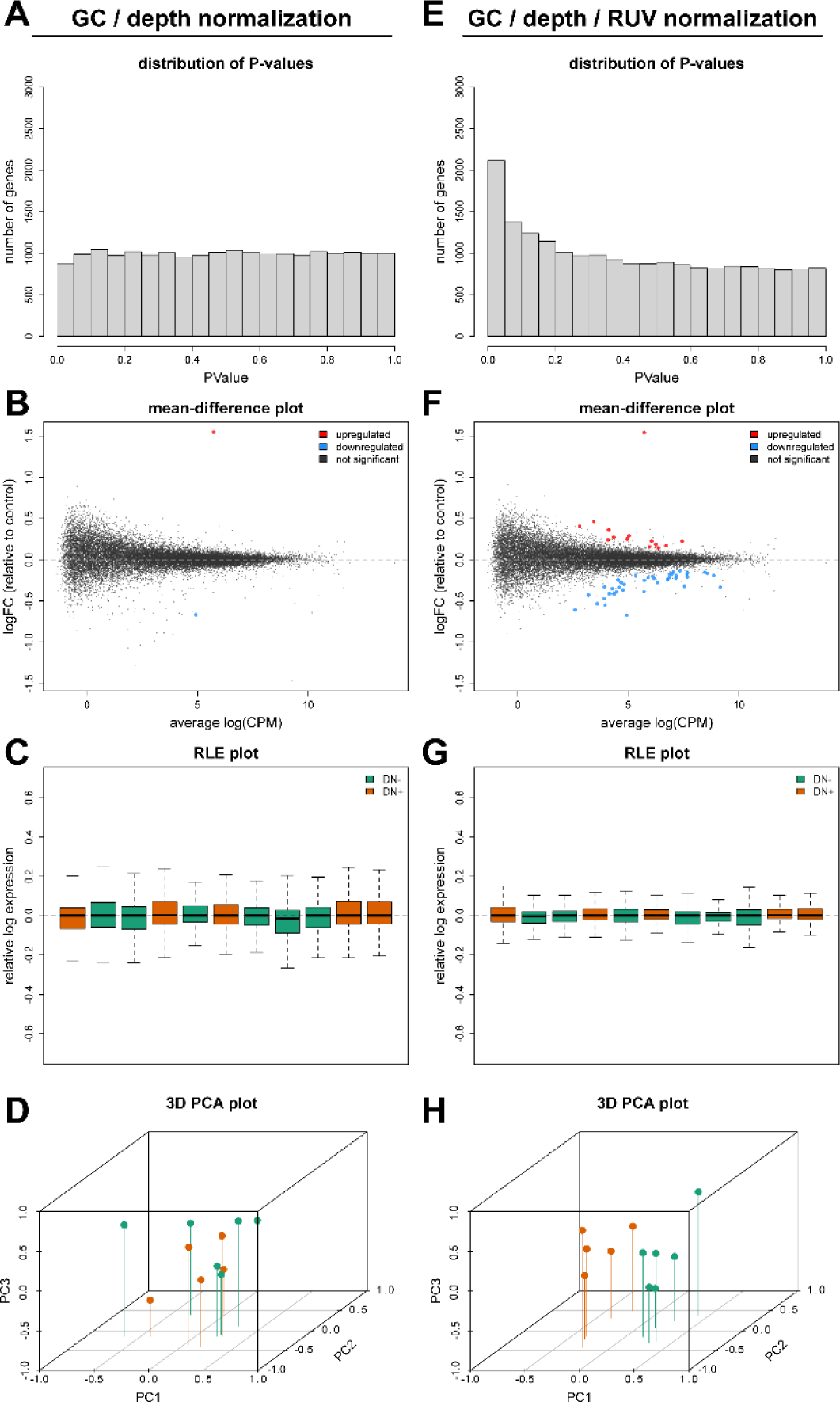
Remove Unwanted Variation (RUV) normalization for analysis of RNAseq after learning in Nr4ADN and control mice. RUV normalization removes unwanted variation that dwarfs biological signal in RNA-sequencing data. **(A-D) Exploratory data analysis without RUV normalization. (A)** Uncorrected P-values from the differential expression analysis in the absence of RUV normalization are uniformly distributed and lack an expected peak at P < 0.05. (**B**) Few differences are statistically significant at a false discovery rate (FDR) of <0.05 after multiple testing correction, despite several genes showing strong fold change trends. (**C**) RLE and (**D**) PCA plots reveal that traditional normalization approaches fail to separate biologically meaningful groups using three principal components. **(E-H) Exploratory data analysis with RUV normalization.** Removing latent sources of variation allows for the separation of experimental groups and increases the power to detect statistically significant differences in gene expression. DN- (Control: tTA+, Nr4ADN-) and DN+ (Nr4ADN: tTA+, Nr4ADN+).

**Fig. S12.**
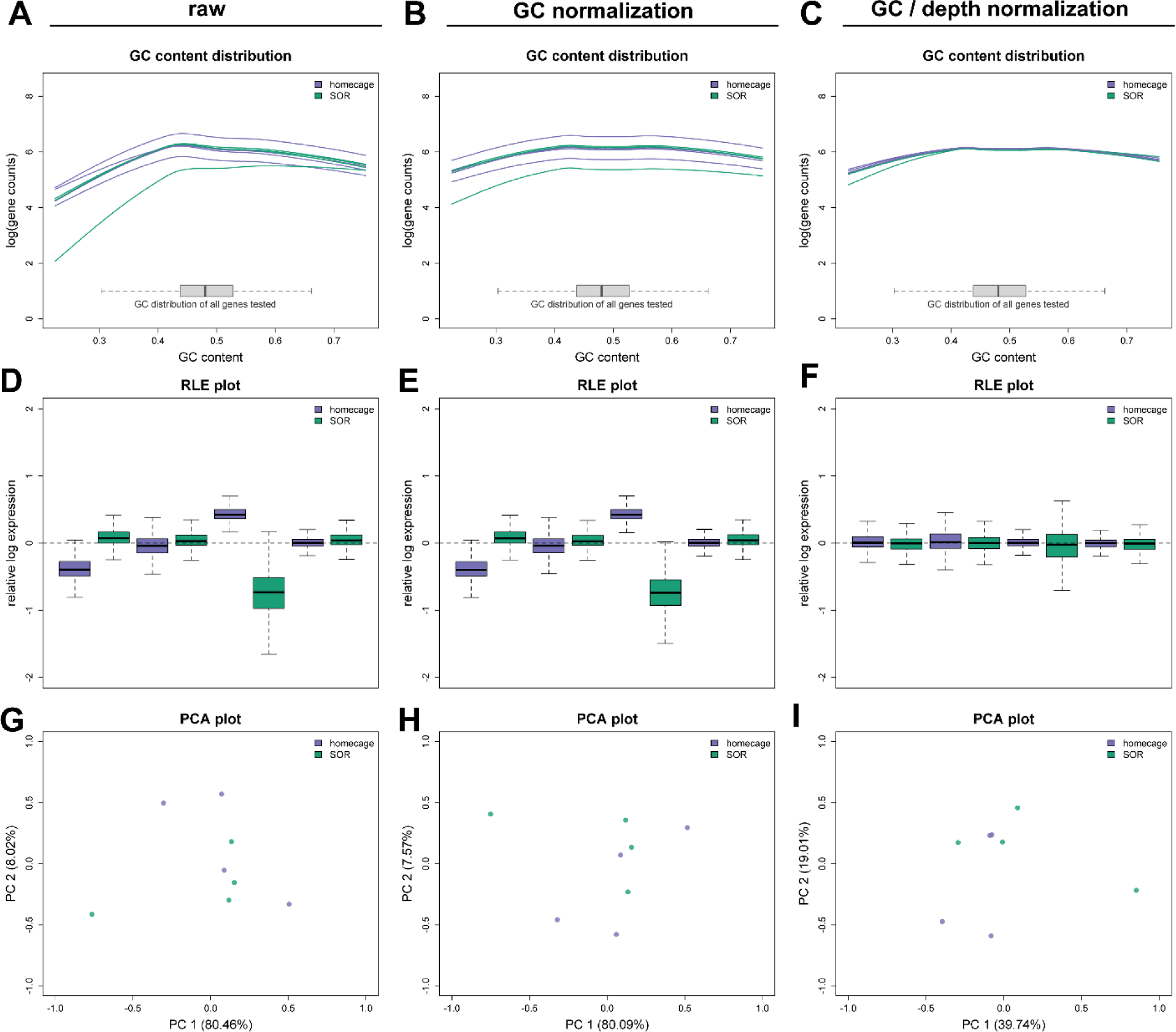
Normalization of differences in GC content distribution and sequencing depth using EDASeq for comparisons of RNAseq studies between homecage control mice and trained control mice. **(A-C)** GC content distributions before normalization (**A**), after full quantile GC content normalization (**B**), followed by upper quartile sequencing depth normalization (**C**). (**D-F)** Relative log expression (RLE) plots before normalization (**D**), after full quantile GC content normalization (**E**), followed by upper quartile sequencing depth normalization (**F**). **(G-I)** Principal component analysis (PCA) plots before normalization (**G**), after full quantile GC content normalization (**H**), followed by upper quartile sequencing depth normalization (**I**).

**Fig. S13.**
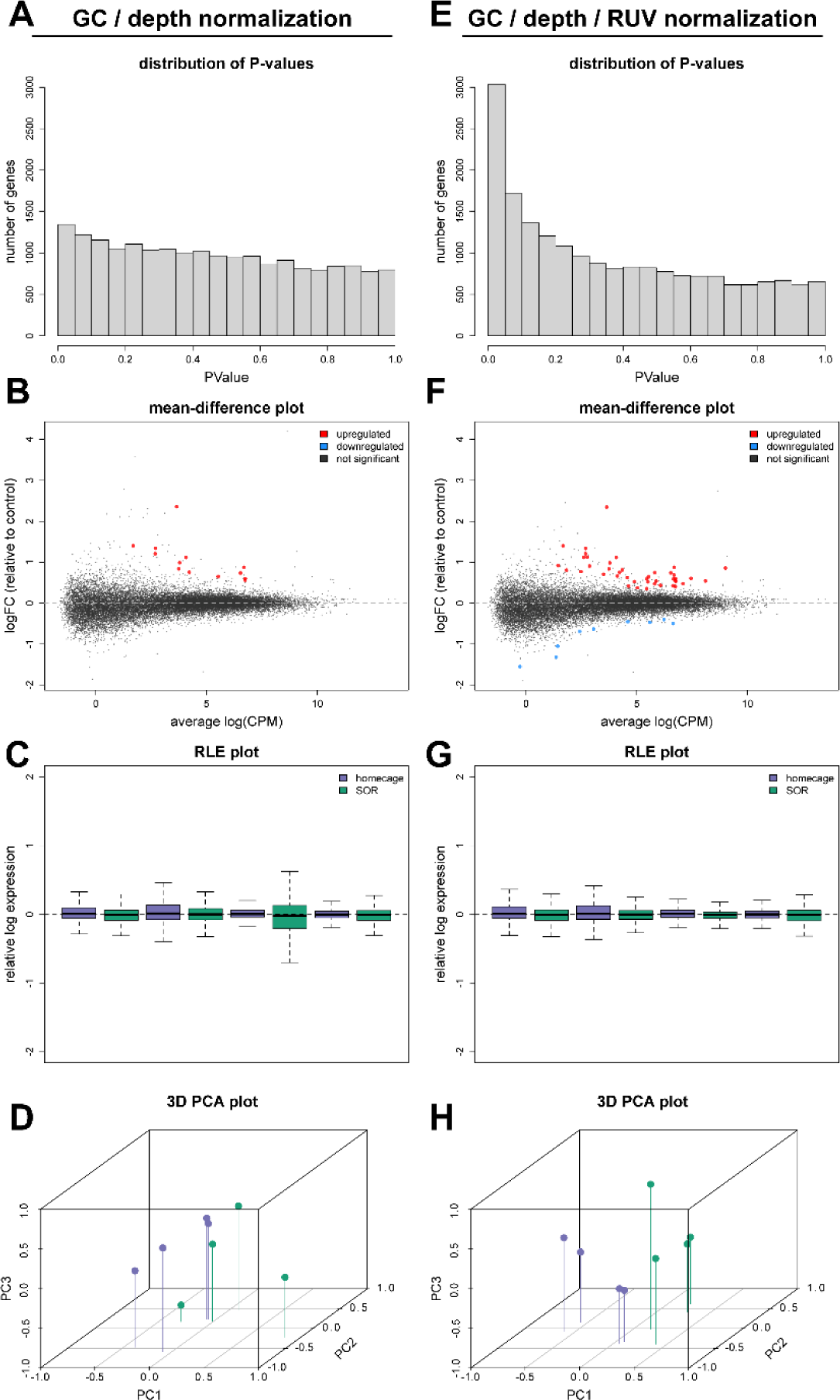
RUV normalization for analysis of RNAseq between homecage control mice and trained control mice. RUV normalization removes unwanted variation that confounds biological signals in RNA-sequencing data. **(A-D), Exploratory data analysis without RUV normalization. (A)** Uncorrected P-values from the differential expression analysis in the absence of RUV normalization are uniformly distributed and lack an expected peak at P < 0.05. (**B**) Few differences are statistically significant at a false discovery rate (FDR) of <0.05 after multiple testing correction, despite several genes showing strong fold change trends. **(C)** RLE and **(D)** PCA plots reveal that traditional normalization approaches fail to separate biologically meaningful groups using three principal components. **(E-H), Exploratory data analysis with RUV normalization.** Removing latent sources of variation allows for the separation of experimental groups and increases the power to detect statistically significant differences in gene expression.

